# Scalable integration and prediction of unpaired single-cell and spatial multi-omics via regularized disentanglement

**DOI:** 10.1101/2025.11.25.689803

**Authors:** Jianle Sun, Chaoqi Liang, Ran Wei, Peng Zheng, Hongliang Yan, Lei Bai, Kun Zhang, Wanli Ouyang, Peng Ye

## Abstract

Deciphering cellular states requires methods capable of integrating large-scale heterogeneous single-cell and spatial omics data. However, these data are typically unpaired due to destructive assays and further confounded by modality heterogeneity, technical noise, and immense scale. Here we present scMRDR, a scalable computational framework based on regularized disentangled representation learning for integrating fully unpaired single-cell and spatial multi-omics datasets. Built on a unified and structure-preserving architecture, scMRDR removes the need for pairing supervision while maintaining computational efficiency, enabling scaling to large datasets spanning multiple disparate omics modalities. Across diverse real-world benchmarks, scMRDR demonstrates strong performance in batch correction, modality alignment, and biological signal preservation. The framework further supports cross-modal translation across omics modalities and enables spatial coordinate imputation for non-spatial single-cell datasets using a reference atlas. The resulting spatial mapping allows spatially resolved analyses, including identification of spatially variable genes and characterization of epigenetic regulatory programs in their native tissue context. These capabilities position scMRDR as a scalable and versatile framework for large-scale multi-omics integration.

## 1 Introduction

Deciphering cellular identity and function requires a holistic picture that captures the complex interplay across distinct molecular layers. The rapid development of sequencing technologies in single-cell and spatial multi-omics has enabled the profiling of diverse molecular characteristics, ranging from transcriptomic landscapes and chromatin accessibility to protein abundance and spatial architecture. However, the destructive nature of most sequencing protocols necessitates measuring different omics data in separate cell populations, resulting in datasets that are completely unpaired [2]. This problem is further exacerbated by the inherent heterogeneity of omics modalities, the pervasive technical noise, and the burgeoning scale of contemporary datasets [3– 5]. Integrating these disparate measurements into a coherent understanding of cellular states remains challenging.

Existing multi-omics integration methods attempt to translate these disparate omics data into a unified cell representation but remain limited in key regimes. Joint dimensionality-reduction frameworks [6] and deep generative models [7] often rely on paired or partially paired observations [8, 9] or prior correspondence information [10, 11], restricting their applicability to unpaired real-world scenarios. Alternative approaches based on manifold alignment [12, 13] or optimal transport (OT) [14, 15] typically require the computation of global cell-cell coupling matrices, which not only restricts to dual-modality integration but also introduces prohibitive computational overhead. As a result, a computational framework that can integrate massive, increasingly complex, and completely unpaired single-cell multi-omics datasets while preserving biological structure remains lacking.

Here, we introduce scMRDR (single-cell Multi-omics Regularized Disentangled Representations), a scalable computational framework designed to integrate completely unpaired single-cell and spatial multi-omics datasets (Fig. 1a). scMRDR leverages a unified *β*-variational autoencoder to disentangle each cell’s latent representation into a modality-shared component that captures shared biological signals, and a modality-specific component that models modality-dependent variation. To ensure that the modality-shared component is well aligned across modalities while remaining biologically faithful, scMRDR couples an adversarial regularization loss for cross-omics alignment with an isomeric regularization loss designed to preserve intrinsic cellular structure. Further, a masked reconstruction objective is designed to facilitate principled learning from the missing or non-overlapping features ubiquitous in multi-omics data. Its unified, structure-preserving architecture allows scMRDR to eliminate the need for pairing information, reduce computational overhead, and scale to massive single-cell and spatial multi-omics datasets.

**Fig. 1.**
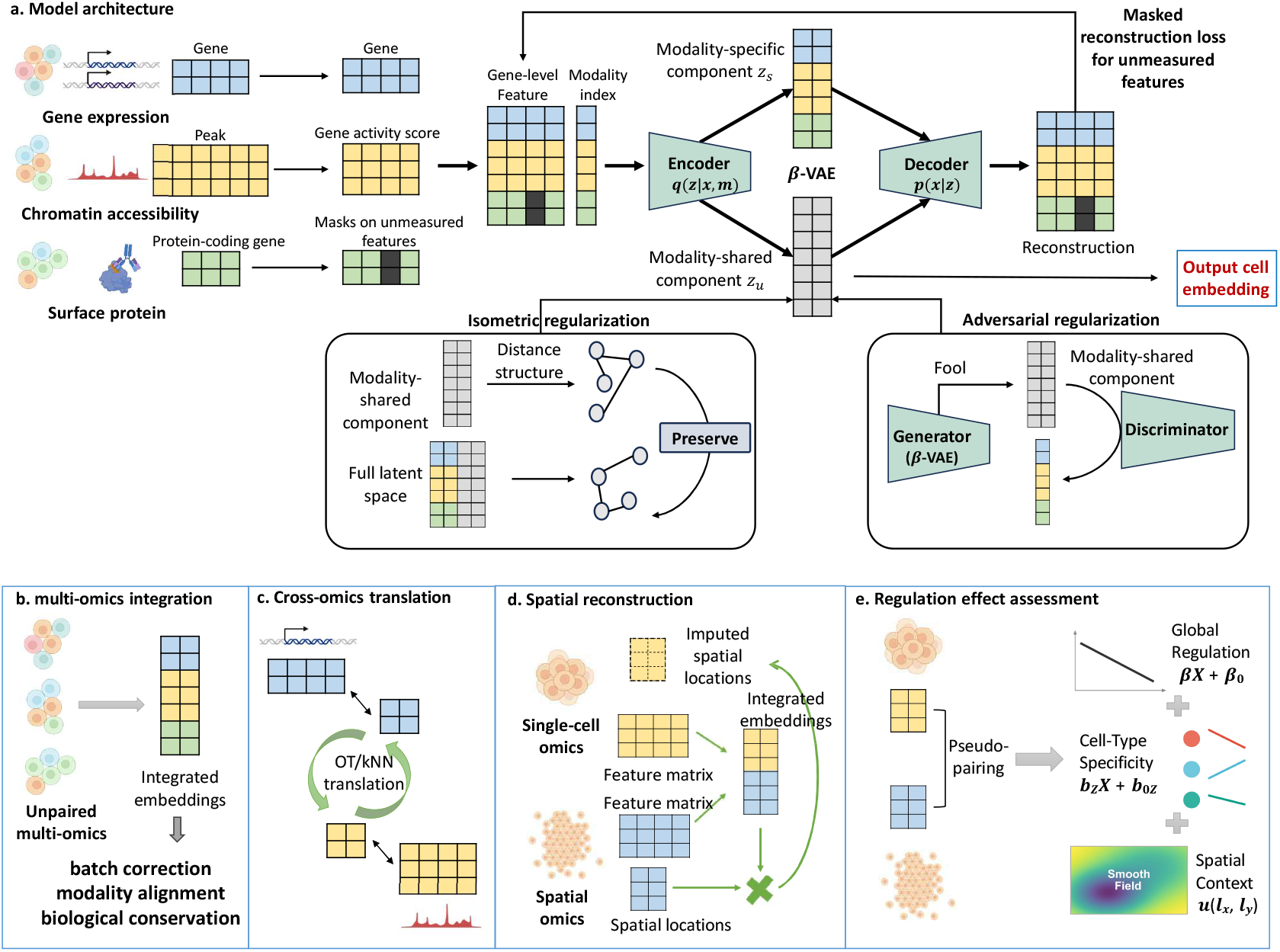
Overview of the proposed scalable computational framework scMRDR, which serves as a powerful tool to integrate unpaired single-cell and spatial multi-omics data. **a**. Model architecture. scMRDR employs a well-designed *β*-VAE to disentangle modality-shared and modality-specific latent components from unpaired multi-omics data. Meanwhile, adversarial regularization and isometric regularization respectively enforce effective modality alignment and biological structure preservation, while masked reconstruction objective mitigates feature mismatch and data sparsity. **b**. Multi-omics integration. By mapping unpaired omics data into a unified latent space while preserving biological signals, scMRDR enables robust batch correction, modality alignment, and biological conservation. **c**. Cross-omics translation. By generating unified and structure-preserving embeddings, scMRDR enables accurate translation across modalities and robust imputation of missing omics features. **d**. Spatial reconstruction. By integrating spatial reference data with non-spatial single-cell profiles, scMRDR accurately infer spatial coordinates for dissociated cells. **e**. Epigenetic regulation analysis. Leveraging spatially resolved multi-omics landscape, scMRDR enables precise characterization of epigenetic regulatory effects through mixed-effects modeling.

Across benchmark datasets with varying scales and modalities, scMRDR shows excellent and robust performance in batch correction, modality alignment, and biological signal preservation (Fig.1b). Besides, it further enables cross-omics translation like predicting chromatin accessibility from gene expression (Fig.1c), and spatial coordinate imputation for dissociated single-cell data with spatial reference atlases (Fig.1d). Crucially, by transforming separate omics layers into pseudo-paired data, scMRDR enables spatially informed downstream analyses, including the identification of spatially variable molecular features and the statistically rigorous dissection of epigenetic regulation effects within native tissue contexts (Fig.1e). In particular, spatial mixedeffects model based on our integrated pseudo-pairs reveals distinct and complementary spatial epigenetic regulatory landscapes, highlighting differential associations of DNA methylation with stable cellular identity and dynamic gene expression programs. Together, scMRDR provides a computational bridge from disparate multi-omics measurements to unified cell representations. As increasingly comprehensive and complex single-cell and spatial multi-omics datasets become available, scMRDR offers a flexible and scalable framework that advances integrative biological discovery.

## 2 Results

### 2.1 Overview of scMRDR for unparied multi-omics integration

To address the challenge of integrating completely unpaired single-cell datasets, we developed scMRDR, a scalable generative framework (Fig.1a). The model is built upon a unified *β*-variational autoencoder (*β*-VAE) that utilizes a single encoder-decoder architecture to process all modalities, which disentangles the cellular state of each sample into a common, modality-invariant latent space (***z***_*u*_) and multiple modality-specific latent spaces (***z***_*s*_). A key innovation of our approach is to ensure that the common ***z***_*u*_ space is both well-aligned across omics and simultaneously preserves the intrinsic biological structure of the data. We achieve this by introducing two specialized regularizations. First, an adversarial regularization loss aligns the ***z***_*u*_ distributions from different omics by training the VAE encoder to “fool” a modality discriminator. Second, an isomeric regularization loss acts as a structure-preserving regularizer, compelling the shared ***z***_*u*_ subspace to retain the topological (e.g., pairwise distance) information present in the full, combined latent space ***z*** = (***z***_*u*_, ***z***_*s*_). Besides, in order to address the sparsity and feature mismatch characteristic of real-world multi-omics data, a masked reconstruction loss is further introduced, with either a gaussian or a ZINB loss function [7] employed according to the distribution of each omics modality. The resulting unified and structure-preserving embeddings ***z***_*u*_ further enable robust downstream analyses, including multi-omics integration (Fig.1b), cross-omics translation (Fig.1c), spatial reconstruction (Fig.1d), and epigenetic regulation analysis (Fig.1e).

### 2.2 scMRDR effectively integrates unpaired scRNA and scATAC data while preserving biological structures

To comprehensively assess the performance of scMRDR, we benchmarked it against nine existing methods: Seurat v5 [16], Harmony [17], scVI [7], scGLUE [10], JAMIE [11], UnionCom [12], Pamona [15], MaxFuse [18], and SIMBA [19], where scVI can be regarded as a baseline counterpart, as it represents a similar generative approach but lacks the disentanglement and regularization components of our model.

The comparison was first performed on a public unpaired dataset of human kidney tissue [20], which includes scRNA-seq data (19,985 cells profiling 27,146 genes) and scATAC-seq data (24,205 cells profiling 99,019 peaks). For scMRDR, scATAC-seq peak signals were aggregated to gene-level activity scores via episcanpy [21] to generate an aligned multi-omics feature matrix. These scores were also used for integration by Seurat, Harmony, and scVI, while the remaining methods utilize raw peak signals as input. The union of highly variable genes identified from each modality served as the input feature space for all methods.

We conducted a detailed evaluation using multiple scib-based metrics [22]. The results demonstrate that scMRDR substantially outperforms the competing methods, showing superior performance in modality integration, biological signal conservation (bio-conservation), and batch correction (Fig.2a, d, Supplementary Fig.S1, S2). Notably, scMRDR produced well-separated embedding clusters that correspond to distinct cell types in an unsupervised manner, confirming the preservation of underlying biological heterogeneity. Simultaneously, samples originating from different omics and technical batches were effectively fused and aligned within the latent space, indicating successful correction of both modality-specific variations and technical noise. In contrast, while several other methods (e.g., Harmony, scVI, and JAMIE) could preserve some biological differences between cell types, they failed to properly integrate the global distributions of the two omics modalities. Systematic ablation studies and sensitivity analyses further verify the function/contribution of each regularization component in the model (Supplementary Table S2).

**Fig. 2.**
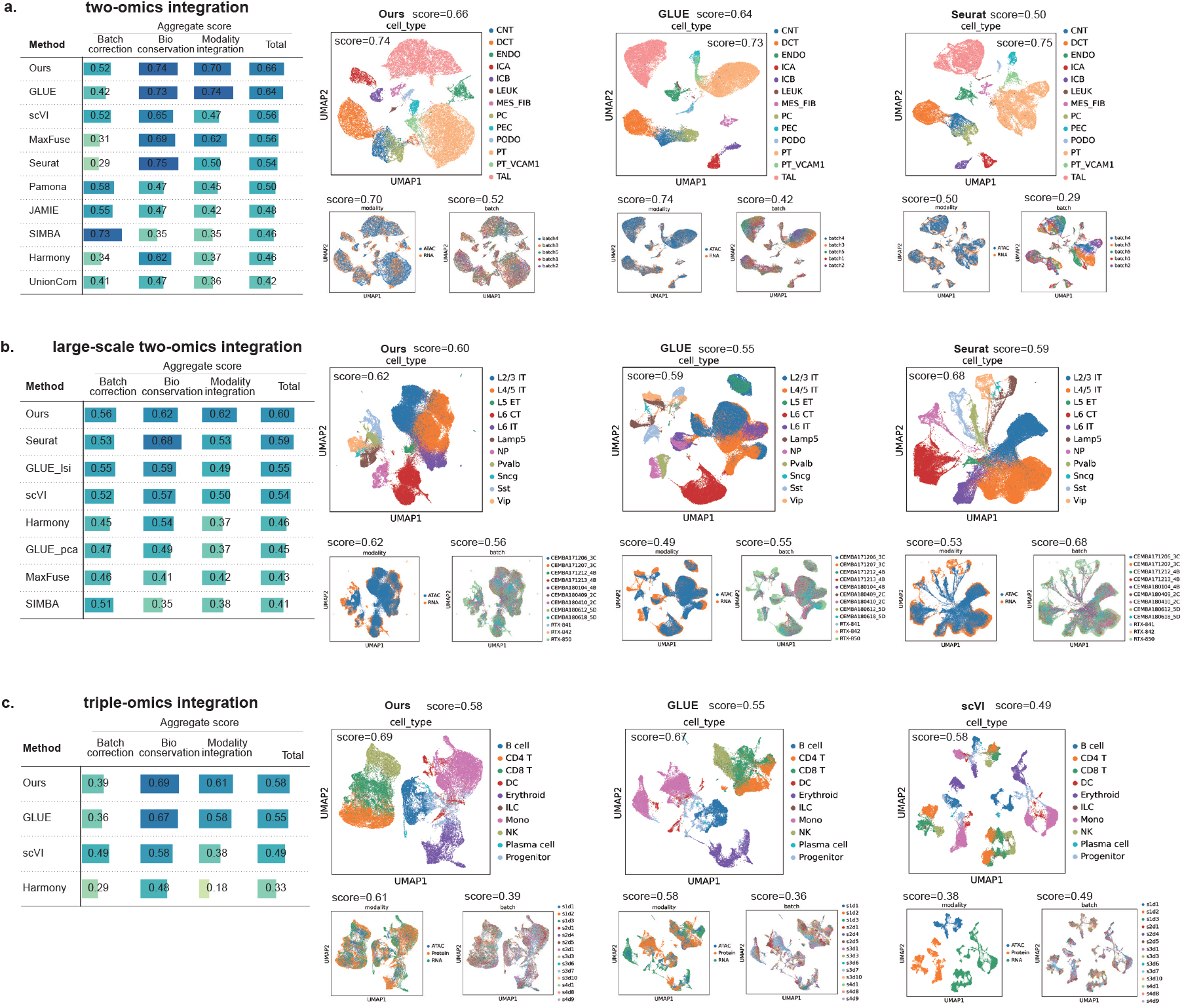
The proposed scMRDR framework achieves robust and scalable integration across increasingly challenging multi-omics integration settings. Unscaled scIB benchmark metrics and UMAP visualizations demonstrate that scMRDR consistently achieves superior or balanced performance in biological conservation (cell-type structure preservation), modality alignment, and batch correction across standard two-omics (**a**), large-scale two-omics (**b**), and challenging triple-omics (**c**) integration tasks. Detailed values of scIB metrics are shown in Supplementary Fig. S1. Visualizations of representative competing methods like GLUE, Seurat, and scVI are shown, while the remaining methods are shown in Supplementary Fig. S2-S4.

### 2.3 scMRDR scales to large datasets and integrates triple-omic profiles

To validate the scalability of scMRDR, we then evaluated its performance on a large-scale dataset from the mouse primary motor cortex, comprising 69,727 cells (27,123 genes) profiled by scRNA-seq and 54,844 cells (148,814 peaks) by scATAC-seq [23]. Notably, several methods based on OT or other global unsupervised manifold alignment (e.g., JAMIE, UnionCom, and Pamona) fail to run on data of this scale due to memory or optimization errors. We therefore compared scMRDR against the remaining methods. Crucially, we observe that some methods that perform adequately on smaller datasets, suffer significant performance drops on this larger dataset (Fig.2a, Supplementary Fig.S1, S3). For example, GLUE’s performance exhibits a strong susceptibility to preprocessing dimension reduction strategies, which are themselves markedly dependent on data scale. In contrast, bypassing the need for explicit preprocessing dimension reduction, scMRDR maintains stable and robust performance, confirming its high scalability for large-level single-cell data integration and its capacity to provide valuable biological insights (Fig.2b, e).

We further demonstrated the remarkable capacity of scMRDR by integrating more than two modalities, a feature not supported by most existing methods. We conducted a case study integrating scRNA-seq, scATAC-seq, and sc-protein levels measured by CITE-seq from a human bone marrow benchmark dataset [24]. This integration involved 30,486 cells (13,431 genes) in transcriptome, 10,330 cells (116,490 peaks) in epigenome, and 18,052 cells (134 surface proteins) in proteome. We applied a masked loss function to accommodate the severe feature missingness inherent in the proteomics measurements, and ablation study confirm its benefits (Supplementary Table S3).

Methods that rely on dual-modality bridge integration, such as Seurat v5, are not suitable for the triple omics integration task and were excluded. We hence benchmarked scMRDR against the remaining methods that support triple-omics integration, including GLUE, scVI, and Harmony. The results show that scMRDR yields a consistently excellent performance in this multi-modal integration task (Fig.2c). Conversely, other methods fail to align well the latent distributions across all three omics, struggling particularly when one modality (proteomics) possessed significantly fewer features than the others (Fig.2c, f, Supplementary Fig.S1, S4).

### 2.4 scMRDR enables accurate cross-omics translation and prediction

The unified and structure-preserving embeddings produced by scMRDR are not an end in themselves, but rather a powerful foundation for critical downstream biological inquiries. For example, the integrated representation enables robust cross-modal translation. By learning a coupling matrix (via OT or kNN) between the latent embeddings of two modalities, scMRDR can accurately impute the profile of one omic from another (e.g., predicting chromatin accessibility from gene expression and verse visa).

We validated the model’s cross-omics translation and prediction capabilities on a paired 10x Multiome [2] (scRNA-seq + scATAC-seq) dataset with 9631 human PBMC cells as the ground truth is available here. We randomly split the entire dataset into two halves: the first half retained only the scRNA-seq measurements, and the second half retained only the scATAC-seq measurements, resulting in an unpaired multi-omics dataset. We used scMRDR to first learn a unified representation for this combined dataset. Subsequently, we conducted OT and weighted KNN on the learned integrated representation, respectively, to further learn the sample coupling between the two omics and predict the missing omics for each sample, which was then compared to the measured ground truth.

scMRDR demonstrates superior cross-omic predictive performance through similar topological distribution between ground truth and predictions (Fig.3a), as well as the high Pearson correlation coefficients (Fig.3b, Supplementary Fig.S5). Specifically, OT surpasses kNN by leveraging global rather than local topological features, resulting in predictions that accurately align with the ground truth distributions, while kNN remains a highly tractable and practical approximation for large-scale applications.

**Fig. 3.**
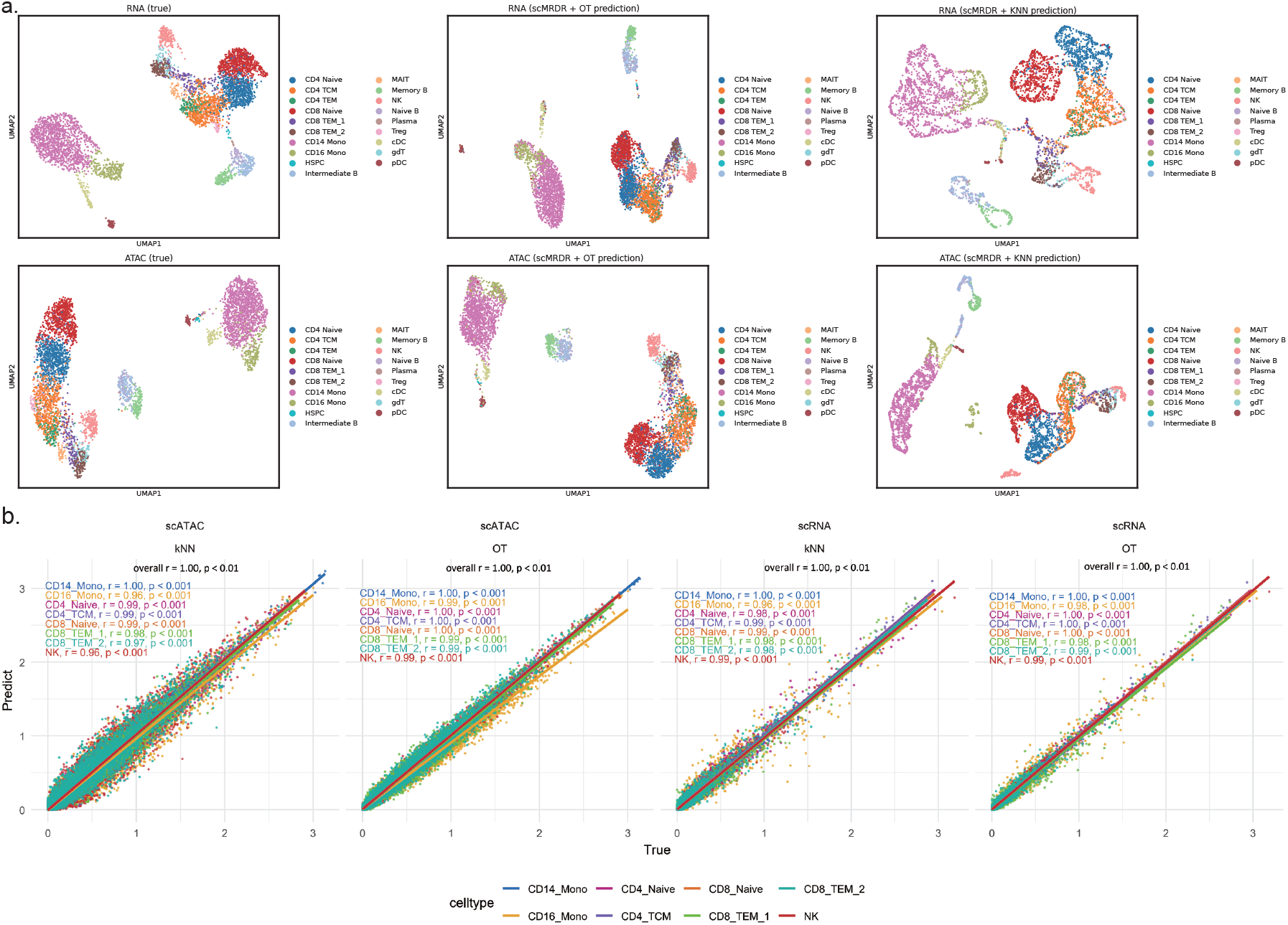
scMRDR enables accurate cross-modality prediction. **a**. UMAP visualizations comparing ground-truth omics profiles with predictions generated by optimal transport (OT) and k-nearest neighbors (kNN). OT-based prediction preserves cell-type topology and global manifold structure, whereas kNN-based prediction introduces structural distortions. **b**. Cell-type-specific quantitative evaluation showing pearson correlation between observed and predicted mean expression levels across genes, demonstrating consistently high accuracy for scMRDR across modalities. Each scatter represents the prediction correlation on one gene.

### 2.5 scMRDR imputes spatial locations for single-cell omics and enhances SVG discovery

Moving beyond single-cell omics to spatial omics, scMRDR demonstrates that its utility is not limited to translating molecular profiles, but also encompasses the prediction of single-cell spatial coordinates. To empirically show that, We integrated scRNA [23], scATAC [23], and spatial transcriptomics (merFISH) [25] of mouse primary motor cortex using our method, and then used the aligned latent representation to interpolate the missing spatial locations in single-cell data by OT. We implemented OT with both entropy-regularized Sinkhorn (Fig.4a) and original EMD algorithms (Supplementary Fig.S6b, S7b), and the spatial visualizations consistently show that this interpolation performs well, where inferred locations of cells align well with the provided cortex layers annotations.

**Fig. 4.**
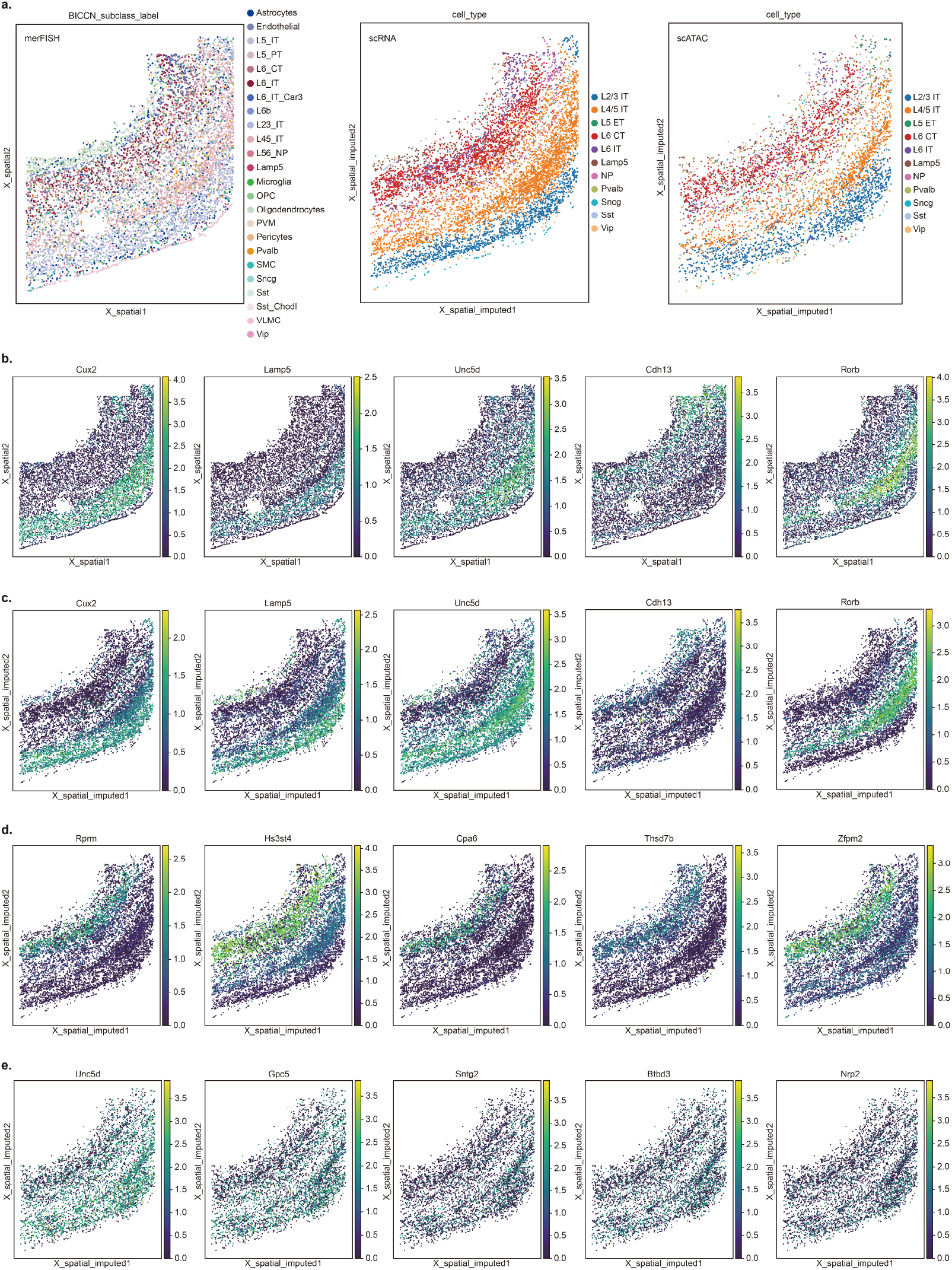
scMRDR integrates spatial transcriptomics with non-spatial multi-omics to reconstruct spatial architecture. **a**. Visualization of the reference merFISH slice alongside scMRDR-imputed spatial mappings of dissociated scRNA-seq and scATAC-seq profiles, colored by cell type, illustrating recovery of large-scale spatial organization. **b**. The top five9conserved Spatially Variable Genes (SVGs) identified in both the original merFISH reference and the imputed scRNA-seq data, visualized in the native merFISH spatial coordinates. **c**. Visualization of the same conserved SVGs on the scMRDR-imputed scRNA-seq spatial map, demonstrating consistent spatial expression patterns between reference and imputed data. **d**. Additional SVGs identified from the scMRDR-imputed scRNA-seq spatial map under the same statistical criteria, highlighting candidate spatial patterns not detected in the reference dataset. **e**. SVGs inferred from gene activity scores in the scMRDR-imputed scATAC-seq spatial map, indicating that spatial variability can be extended to chromatin accessibility-derived signals. Together, these results show that scMRDR enables spatial mapping of non-spatial single-cell multiomics data and supports downstream spatially informed analyses, including the characterization of spatially variable molecular programs.

Due to the low coverage of merFISH (only 254 genes measured), only 103 genes are detected as spatial variable genes (SVGs) by SPARK-X [26] (*P*_adj_ *<* 10^−20^). We leveraged the spatially interpolated scRNA data (26069 genes) and 4095 SVGs (*P*_adj_ *<* 10^−20^) are detected. We replicated 83 out of 101 SVGs detectable by merFISH (like *Cux2, Lamb5, Unc5d* ) (Fig.4c, d), and also revealed new SVGs (like *Hs3st4, Cpa6, Zfpm2* ) (Fig.4d, Supplementary Fig.S9). Similarly, using scATAC with imputed spatial locations, we identified 142 SVGs in gene activity scores (*P*_adj_ *<* 10^−20^), including several key transcription factors like *Unc5d, Gpc5, Sntg2, Nrp2* (Fig.4e, Supplementary Fig.S10). This will further support the investigation of spatially specific regulatory mechanisms.

For example, the *Unc5d* gene, encoding a member of the UNC-5 family of netrin receptors, demonstrates a highly specific spatial pattern, strongly concentrated in Layer 4 (L4), specific primary sensory areas of the neocortex including primary somatosensory cortex (S1), the primary visual cortex (V1), and the primary auditory cortex (A1), which suggests its crucial role in the development and maturation of sensory cortical circuits [27]. The *Zfpm2* gene (also known as *Fog2* ) exhibits a highly specific spatial distribution in the mouse cerebral cortex, primarily functioning as a key molecular marker for Layer 6. Its expression is largely restricted to the deepest cortical layer, where it is essential for the proper differentiation of corticothalamic projection neurons (CThPNs), playing a particularly critical role in regulating the identity and axonal projections of CThPN subsets within the motor cortex [28].

### 2.6 scMRDR integrates structurally heterogeneous omics to elucidate the spatial epigenetic regulation landscape via mixed-effects modeling

To further demonstrate scMRDR’s performance in integrating multi-omics data with distinct resolutions and more heterogeneous data structures, we integrated gene expressions (scRNA-seq), DNA methylation ratios measured by single-nucleus methylation sequencing (snmC-seq) on both CG (mCG) and non-CG (mCH) sites [29], and spot-resolution 10x Visium spatial transcriptomics data from mouse frontal cortex. The 10x Visium spots serve as a spatial reference for interpolating single-cell RNA expression and DNA methylation ratio data. Our spatial interpolation results accurately map different cell types to their corresponding cortical layers, similar to the effect achieved by SIMO [30]. However, while SIMO requires multi-step integration and explicitly utilizes pre-acquired cell type labels, we achieve a completely unsupervised, unified integration of multiple modalities, adapted to the vast and increasingly ubiquitous volumes of unannotated data. This is particularly crucial for emerging omics modalities lacking sufficient reference atlases, where their annotations typically need to be derived through the unsupervised alignment with well-characterized reference modalities.

This spatially resolved integration of transcriptomic and epigenomic data further empowers the investigation of regulatory effects underlying the spatial heterogeneity of gene expression. We aggregated and aligned the spatially imputed scRNA-seq and snmC-seq data according to their spatial coordinates to generate pseudo-paired multiome data. Leveraging a Spatial Generalized Additive Mixed Model (SGAMM) to account for spatial autocorrelation and cell-type heterogeneity, we dissected the epigenetic regulation of gene expression in the mouse prefrontal cortex (Fig.5b-e). Our analysis reveals a pervasive repressive landscape, where DNA methylation, particularly in the non-CG context (mCH), shows a robust negative association with gene expression. This transcriptional silencing effect is most pronounced in canonical interneuron markers such as *Sst, Pvalb*, and *Dlx5*, aligning with recent single-cell methylome atlases which establish mCH accumulation as a critical hallmark of neuronal maturation and lineage specification [23, 31]. The strong signal detected in the mixed model underscores the role of mCH not merely as a transient regulator, but as a stable epigenetic mechanism that enforces cellular identity by silencing lineage-inappropriate genes.

**Fig. 5.**
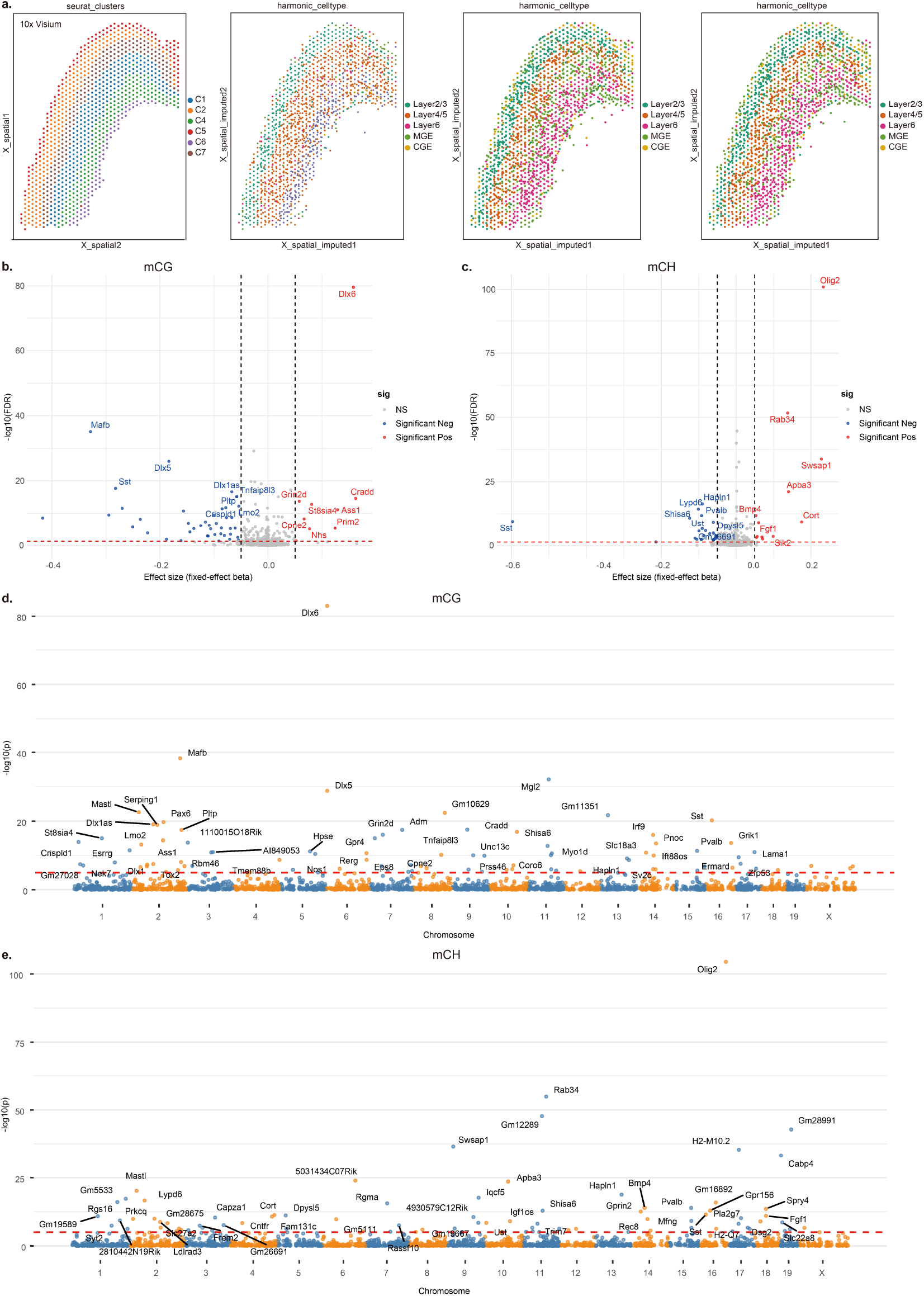
scMRDR integrates spot-base spatial omics and single-cell data for spatial reconstruction and mechanistic dissection of the epigenetic landscape. **a**. Visualization of the reference 10x visium spots and scMRDR-imputed spatial distributions of scRNA-seq, snmC-mCG, and snmC-mCH profiles, which reveals consistent laminar and regional organization across modalities. **b-c**. Identification of spatially associated epigenetic signals using Spatial Generalized Additive Mixed Model (SGAMM). Volcano plots show the fixed-effect coefficients (*β*) and corresponding statistical significance (-log10 FDR) for mCG (b) and mCH (c) sites, after controlling for cell-type composition and spatial auto-correlation. **d-e**. Genome-wide landscapes of spatially associated epigenetic regulation. Manhattan plots summarize spatial association statistics across chromosomes for mCG (d) and mCH (e), with prominent peaks corresponding to genes with significant epigenetic regulation effect, where the red dashed line indicates the significance threshold *P* = 1*e* − 5 since thousands of genes are included.

Notably, comparing these results with a cell-type fixed-effect model (SGAM, Supplementary Fig.S11) provides critical mechanistic nuance. For instance, the strong signal observed for *Olig2* in the mixed model (Fig.5b) was effectively ablated in the fixed-effect model (Supplementary Fig.S11). This indicates that the *Olig2* signal is largely driven by broad compositional differences between cell types (i.e., the stark contrast in methylation profiles between oligodendrocytes and neurons) rather than intra-cell-type regulation. The attenuation of such signals in the fixed-effect model is consistent with the interpretation that the strong methylation-expression coupling observed globally is substantially driven by compositional differences between cell types rather than dynamic intra-cell-type tuning. This distinction is consistent with the “genomic barrier” hypothesis, where methylation (particularly mCH) landscapes may function as a stable marker of cellular identity established during development to restrict cell fate, rather than acting as rapid transcriptional switches in the mature brain [29, 32].

### 2.7 scMRDR enables efficient cross-species atlas-scale integration

Finally, to further demonstrate the feasibility of scMRDR on atlas-scale data, we integrated a cross-species scRNA dataset comprising a total of 610,718 brain cells from primary frontal cortex of human, macaque, marmoset, and chimpanzee [33], where gene names of homologous genes have been harmonized. Similar to experiments above, the integration was performed in a fully unsupervised manner without using any explicit cell-type annotations. The results (Supplementary Fig.S13) show that similar cells across different species are well integrated at the level of cell subtypes, demonstrating the model’s excellent ability to achieve accurate and biologically meaningful cross-species integration. Beyond showing its scalability to massive atlas, this also suggests that scMRDR can serve as a robust and powerful framework for annotating reference-poor species and decoding the evolutionary conservation of specific cell types.

## 3 Discussion

Translating highly disparate multi-omics data into a holistic cellular picture is essential in understanding the complex regulatory networks that govern cellular identity, function, and spatial organization. In this study, we introduce scMRDR, a scalable generative framework for integrating multiple completely unpaired single-cell omics datasets. A fundamental challenge in unpaired integration lies in aligning modalities robustly while preserving fine-grained biological structure, such as rare cell states and subtle subtype variation. Many existing approaches depend on explicit pairing information [8, 9], which is often unavailable when datasets are generated independently, or rely on computationally intensive global coupling strategies that scale poorly in large atlas-level settings [12–15]. Several recent studies [34–36] have begun exploring latent-space decomposition strategies for integration. However, many such approaches still rely on partial pairing supervision, stacked modality-specific encoders, or cross-modal contrastive objectives, which can limit scalability and flexibility in fully unpaired, multi-omics scenarios.

In contrast, scMRDR addresses these challenges through a unified *β*-VAE architecture equipped with a dual-regularization scheme. Specifically, scMRDR adopts a single encoder-decoder architecture under a unified *β*-VAE framework, treating observations from different omics layers as instances within a shared generative model. Additionally, an adversarial objective is used to align the shared latent representations (***z***_*u*_) across modalities, while an isometric regularization term constrains this shared space to preserve the intrinsic topological structure encoded in the full latent space. Together with a masked reconstruction loss that accommodates non-overlapping feature spaces, this design enables robust cross-modal alignment without requiring paired cells or explicit global cell-cell correspondence maps, making it a flexible and scalable framework across completely unpaired datasets spanning multiple omics modalities.

The utility of this integrated embedding is demonstrated in two key downstream tasks: cross-modal imputation and spatial coordinate imputation. The latter is particularly impactful. Firstly, imputing the spatial location of scRNA-seq data will greatly alleviate the limitations of traditional spatial transcriptomics methods in terms of resolution (e.g., Visium) or sequencing coverage (e.g., MERFISH). This makes it possible to quantify the spatial expression of a larger number of genes at single-cell resolution, benefiting further spatial analysis like SVGs detection. Secondly, and more importantly, this imputation method effectively expands the multi-omics horizon of spatial genomics for omics types where spatial location is currently difficult to measure, such as scATAC-seq and single-cell methylation. This allows for the spatial heterogeneity of more transcriptional regulatory processes to be revealed.

We showed that by imputing spatial locations for non-spatial scRNA-seq and snmc-seq data, scMRDR enables the application of advanced statistical models such as SGAMM, incorporating cell-type-specific effects and confoundings like spatial auto-correlations. This specific application highlights a critical advance: the ability to move beyond simple data harmonization to a framework that allows for rigorous statistical testing of biological hypotheses (e.g., the effect of DNA methylation on mRNA expression). This approach explicitly corrects for complex confounders, such as continuous spatial autocorrelation and cell-type-specific heterogeneity, which are often ignored in standard analyses.

Despite its excellent performance, scMRDR has some limitations. First, although the disentanglement between ***z***_*u*_ and ***z***_*s*_ is encouraged by the *β*-VAE objective and additional regularization terms, it is not theoretically guaranteed to be exact. Second, the isometric loss treats the full latent representation ***z*** learned by the VAE as a surrogate for the “true” underlying data structure; however, this structure is itself model-dependent and may introduce additional bias and unstable posterior. Finally, the selection of hyperparameters for different loss components (e.g., *β, λ*, and *γ*) may require careful tuning, particularly when integrating highly heterogeneous datasets.

Looking forward, extending the framework to accommodate more diverse data modalities represents an important avenue for future research. In particular, integrating heterogeneous formats, such as gene 3D structures, genetic association signals (e.g. GWAS and eQTL) [37] or cell-cell interaction graphs [38] with conventional single-cell omics matrices would substantially broaden the scope of cross-modal biological discovery. n addition, systematic characterization of cellular responses to chemical perturbations and genetic interventions across multiple omics layers [39, 40] calls for principled and flexible methodologies that can align disparate modalities while preserving perturbation-specific heterogeneity.

In summary, scMRDR offers a flexible, scalable and robust solution for unpaired multi-omics integration. By constructing a unified and structure-preserving latent space, it provides a foundation for a wide range of downstream analyses, promoting to bridge large-scale data integration with spatially informed and mechanistically grounded modeling of gene regulatory programs.

## 4 Methods

### 4.1 scMRDR for unpaired multi-omics integration

#### Key idea

We aim to achieve the latent space disentanglement with a generative model designed for single-cell multi-omics. We implement a unified *β*-VAE composed of a single encoder-decoder, treating observations in different omics equally as a single sample, thereby ensuring flexibility and scalability in completely unpaired data across multiple omics. Theoratically, such disentangled subspaces are unidentifiable (i.e., not unique) without additional constraints [41]. We leverage this unidentifiability and, by imposing isometric and adversarial regularization, constrain the modality-shared subspace to be the one that preserves the maximum sample structure information from the entire space while aligning different modalities.

#### Generative model for multi-omics data with disentangled latent space

To achieve flexible and scalable integration, we propose a generative model tailored for single-cell multi-omics data (Fig.1a). We assume that observations ***x***^(*m*)^ in the omics *m* are generated from latent embeddings lying in two independent subspaces, i.e., common latent variables ***z***_*u*_ shared across modalities and modality-specific latent variables 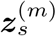 distributional distinctly for different modalities. Then we have

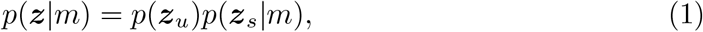

and

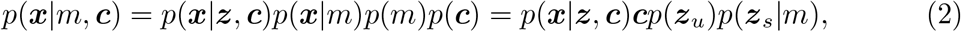

where ***c*** represents other covariates. In particular, batch effects are systematic variations introduced by non-biological factors, including differences in experimental runs, reagent lots, or operators. These effects can obscure true biological signals or introduce spurious patterns in the data [5]. To mitigate such confounding influences, batch information is commonly included as a covariate in the modeling process.

Thanks to the central dogma, sequencing reads of mRNA, protein, epigenomic features, and other omics data can all be mapped to the gene level, such as gene activity scores derived from peak aggregation in scATAC-seq, and methylation scores by averaging ratios across all methylation sites within a gene body. For sparse count data with over-dispersion (i.e., the variance exceeds the mean) and dropouts, we parametrize the generative process *p*(***x*** | ***z, c***) using a zero-inflated negative binomial (ZINB) distribution [7, 42], i.e.,

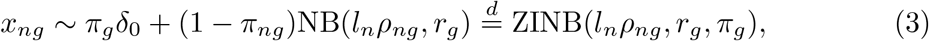

where *δ*_0_ is point mass at zero, *l*_*n*_ represents the library size of cell *n, ρ*_*ng*_ is the mean proportion of the corresponding measurement (RNA expressions, activity score, protein level, etc.) of gene *g* in cell *n, r*_*g*_ is the dispersion factor of gene *g, π*_*ng*_ is the dropout rate of gene *g* in cell *n*. Inspired by scVI [7], we parametrize the non-linear neural networks as follows ***h*** = *f*_*h*_(***z, c***), *ρ*_*ng*_ = *f*_*ρ*_(***h***), *π*_*ng*_ = *f*_*π*_(***h***). When normalized profiles or z-scores or more non-count data (like DNA methylation ratios) included, we can change to a simple Gaussian decoder with 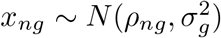.

We employ *β*-VAE [43], an variant of classcial variational autoencoders (VAEs) [44] by introducing a hyperparameter *β* to upweight the KL divergence term in the VAE objective, to model the generative process and encourage disentanglement. Prior distributions of latent factors ***z*** are assumed as isotropic multivariate Gaussian distributions, i.e., 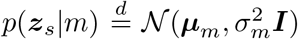 and 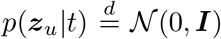. We employ variational posteriors *q*(***z***_*u*_|***x***) = *µ*_*u*_(***x***) + *σ*_*u*_(***x***) ⊙𝒩 (0, ***I***) and *q*(***z***_*s*_|***x***, *m*) = *µ*_*s*_(***x***, *m*) + *σ*_*s*_(***x***, *m*) ⊙ 𝒩 (0, ***I***) to approximate the prior, and the loss function (negative ELBO) of *β*-VAE is

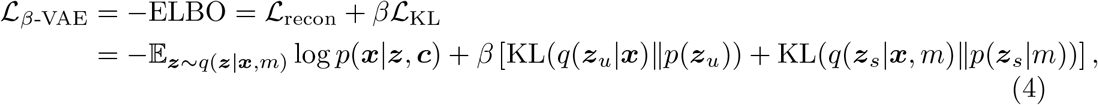

where *β >* 1 to encourage the disentanglement of ***z***_*u*_ and ***z***_*s*_. For the ZINB model, the reconstruction loss, i.e., the expected log likelihood under the variational posterior, is

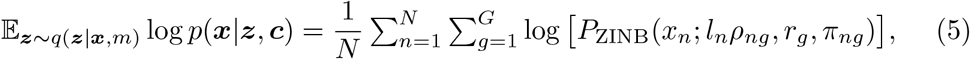

where *P*_ZINB_(*X* = *x*; *µ, r, π*) = *π*𝕀_*x*=0_ + (1 − *π*)*P*_NB_(*x*; *µ, r*), and *P*_NB_(*x*; *µ, r*) stands for the probability mass of negative binomial distribution *NB*(*µ, r*) at *x*, i.e.,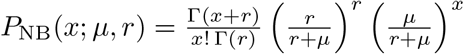.

#### Adversarial regularization for omics integration

To further encourage the alignment of distributions of 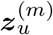 from different omics, we impose an additional adversarial regularities [45, 46] by introducing a *m*-class discriminator 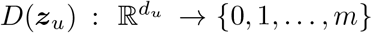 to distinguish ***z***_*u*_ of samples from different omics and try to optimize its capacity, i.e.,

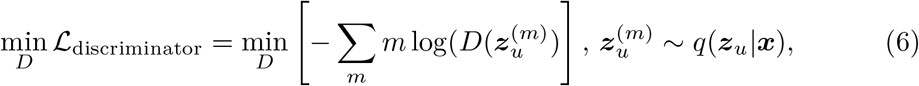

while training the VAE encoders *q*(***z***_*u*_ | ***x***) to fool the discriminator as much as possible by optimizing in the opposite direction

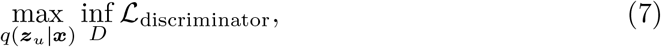

which is equivalent to

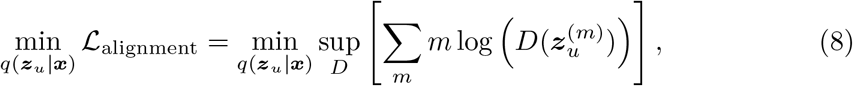

achieving a proper alignment of embeddings from different omics ultimately.

#### Isometric loss for structure preservation

To ensure that ***z***_*u*_ captures the biological heterogeneity among cell populations (e.g., cell types and cell states), we introduce an additional unsupervised structure-preserving regularization, since true cell-type labels or annotations are typically unavailable. Though the original feature matrices are high-dimensional, the full latent representation ***z*** = (***z***_*u*_, ***z***_*s*_) learned by the generative model effectively preserves intra-modality structure. Hence, we reformulate the problem as encouraging ***z***_*u*_ to retain the structural information of the full latent space. Specifically, since the latent embeddings already reside in a low-dimensional space, we apply an isometric loss [47] that minimizes the discrepancy between the pairwise Euclidean distance matrices computed from ***z*** and from ***z***_*u*_, for each modality, i.e., minimizing the Frobenius norm of the distance difference matrix

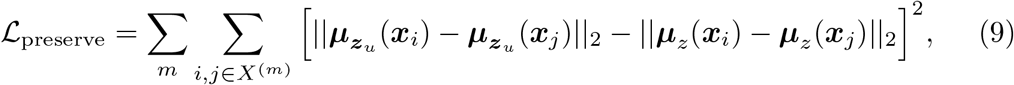

where 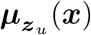 is the posterior mean of variational approximation *q*(***z***_*u*_|***x***) and ***µ***_*z*_ (***x***) is the posterior mean of total latent embeddings (*q*(***z***_*u*_ ***x***), *q*(***z***_*s*_ ***x***, *m*)).

And the total optimization goal becomes

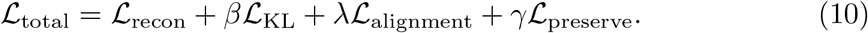

We first update the discriminator by optimizing ℒ_discriminator_, then update VAE with respect to the total loss ℒ_total_ in turn in each training mini-batch.

#### Masked reconstruction loss for features mismatch and missing

Since different modalities are measured disparately, although it is possible to align features across modalities at the gene level, there still exists severe missing and non-overlapping features ubiquitous due to different sequencing coverages. For example, antibody-based protein profiling techniques (like CITE-seq) and *in situ* imaging-based spatial quantification methods (like smFISH) typically covers only a few hundred proteins due to the limited availability of antibody markers [3] or pre-defined fluorescence panel and optical crowding [4], while tens of thousands of genes can be measured in other sequencing-based omics. Restricting the analysis to the overlapping features across all modalities would lead to substantial information loss, whereas naively imputing unmeasured features with zeros would severely distort the data distribution. To address this, we introduce a binary mask ***b*** ∈ {0, 1}^*G*^ indicating feature availability that prevents gradients from back-propagating through unmeasured features in the reconstruction loss for each modality, and then scale by the proportion of available features to ensure that the reconstruction loss for each sample is on a comparable scale, i.e.,

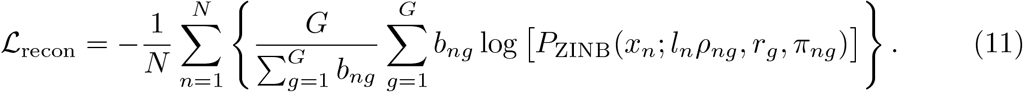

The masked loss strategy ensures that the model can fully utilize the available information while preserving the integrity of the original data distribution.

#### Weighted sampling with severely unbalanced omics samples

Integrating heterogeneous omics faces significant challenges regarding sample size imbalance, especially between large-scale single-cell datasets and single-slice spatial references. To mitigate this disparity, we employ inverse-frequency weighted sampling within each training batch to balance the modalities and stable the training loss. For validation, we maintain the original traversal method to preserve the data’s intrinsic distribution and ensure optimization stability.

### 4.2 Prediction of missing omics using integrated latent embeddings

The obtained integrated latent embeddings were utilized to interpolate missing data across omics modalities and hence enable cross-omics translation. Specifically, given the unified latent embeddings 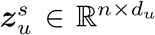 and 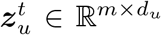 derived from unpaired omics *s* (e.g., scRNA) and omics *t* (e.g., scATAC), we computed a coupling matrix *W* ∈ ℝ^*n*×*m*^ capturing the correspondence between cells in the two datasets. This matrix enables cross-modal prediction, for instance, from *s* to *t*, according to 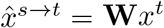. This coupling matrix **W** can be derived using various approaches, such as by solving an OT problem between the 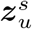 and 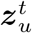 distributions, or by applying kNN algorithm with either hard or soft assignments.

### 4.3 Spatial location imputation of single-cell omics

The scMRDR framework can be further extended to integrate spatial omics data (e.g., spatial transcriptomics) with other non-spatial single-cell modalities. This integration facilitates the imputation of spatial coordinates for the non-spatial single-cell data by leveraging the resulting unified latent embeddings. Specifically, given a spatial omics dataset *s* with known coordinates 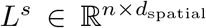 and a non-spatial single-cell dataset *t* (containing *m* cells), we first compute their respective unified latent embeddings, 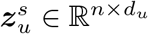 and 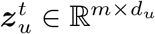. An OT problem is then solved to find the coupling matrix Γ ∈ ℝ^*m*×*n*^ that minimizes the transport cost between the two latent distributions

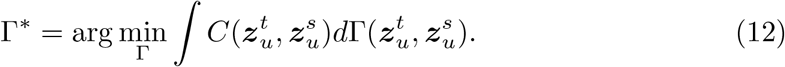

Subsequently, the spatial locations for the cells in dataset 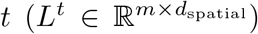 are imputed via barycentric projection. This is achieved by projecting the known spatial coordinates *L*^*s*^ onto dataset *t* using the optimal coupling matrix Γ^∗^, i.e., the barycentric projection

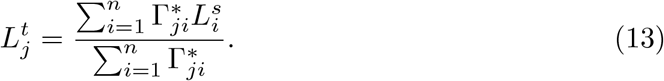

Spatial imputation will alleviate the limitations of traditional spatial transcriptomics methods in terms of resolution (e.g., Visium) or sequencing coverage (e.g., MERFISH), enabling quantification of the spatial heterogeneity of a larger number of genes at single-cell resolution within complex regularization process, for example, the detection of spatial highly variable genes (SVGs).

### 4.4 Test on the effect of DNA methylation on gene expression with spatial generalized additive mixed-effect model

#### Spatial location imputation of scRNA and snmC-seq

We used scMRDR to integrate 10x Visium data of the mouse brain cortex with spatial location measurements and scRNA and snmC-seq data without spatial coordinate information. We used normalized *z*-scores of mRNA expression and DNA methylation levels as input to align RNA and methylation features. Using the unified embedding obtained from the integration, we interpolated the spatial location information of scRNA and snmC-seq cells.

#### Spatial Data Alignment

Next, to create spatially comparable units, we performed a data aggregation step. For both the RNA and methylation datasets independently, we grouped spots that shared the same anatomical layer and occupied a similar spatial location (defined by rounding coordinates to a precision of two decimal places). The gene profiles within each of these unique spatial-anatomical groups were then averaged, creating a new, aggregated “pseudo-spot” dataset for each modality.

Finally, we aligned these aggregated datasets. The matching was performed strictly within each anatomical layer. For every aggregated RNA pseudo-spot, we identified its single nearest-neighboring aggregated methylation pseudo-spot based on Euclidean distance. This process generated the final, one-to-one matched dataset of mRNA expression and DNA methylation profiles used for all downstream analyses.

#### Statistical Analysis with SGAMM

To assess the relationship between gene body DNA methylation status and corresponding mRNA expression level while correcting for spatial confounding and hierarchical cell-type structure, we implemented a Spatial Generalized Additive Mixed-effect Model (SGAMM) for each gene. For each gene, we modeled log-normalized expression (*E*_*i*_) as a function of normalized methylation ratio (*M*_*i*_, computed by averaging mCG and mCH level across the gene body, i.e., TSS to TES) for each cell *i*, located at spatial coordinates *z*_*i*_ = (*x*_*i*_, *y*_*i*_) and belonging to a specific cell type *k*_*i*_.

To account for spatial autocorrelation, we introduce a zero-mean Gaussian Process, *s*(***z***) ∼ 𝒢𝒫(0, **K**) to model the spatial effect at cell coordinates ***z***_*i*_ = (*x*_*i*_, *y*_*i*_). The Gaussian Process is characterized by a covariance function *K*(***z***_*i*_, ***z***_*j*_), which defines the spatial covariance between any two cells *i* and *j* as a function of their locations. This term flexibly captures continuous, spatially-structured variation that is not accounted for by other model components. We also incorporate the cell-type-specific effect and consider two variants.

- Model 1 (SGAM): We implement a Spatial Generalized Additive Model (SGAM) using the mgcv package in R by introducing an interaction term and treating cell type as fixed effects.

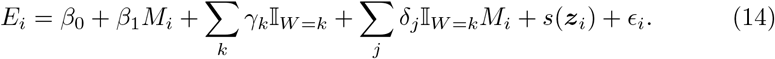
- Model 2 (SGAMM): We implement Spatial Generative Additive Mixed-effect Models (SGAMM) using the gamm4 package in R and use random effects 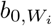 and 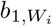 to capture deviations specific to the cell type *W*_*i*_, which are assumed to follow a zero-mean multivariate normal distribution 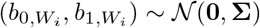.

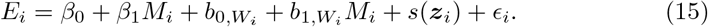

Finally, *ϵ*_*i*_ ∼ 𝒩(0, *σ*^2^) is the independent, non-spatial residual error term. Statistical significance of the methylation association (*β*_1_) was assessed via t-statistics from the fitted model.

### 4.5 Experiments setting and evaluation metrics

We compared to existing methods through a series of experiments on real-world datasets, beginning with standard benchmarks and then scaling up to more complex single-cell and multi-omics scenarios, and evaluate in terms of cell-type clustering, modality integration, and batch removal. Cell-type labels in different omics has been harmonized in evaluation. It should be emphasized that we did not use cell-type labels during training. UMAP visualizations are presented for qualitative comparison. For quantitative evaluation, the following commonly used metrics are included: F1 isolated label scores, k-means NMI, k-means ARI, cell-type Silhouette, and cell-type separation LISI (cLISI) to evaluate the performance in cell type conservation, modality Silhouette, modality integration LISI (iLISI-modality), kBET, Principal Component Regression (PCR) *R*^2^, and graph connectivity to evaluate the performance in modality integration, as well as batch Silhouette, batch integration LISI (iLISI-batch), kBET, and PCR *R*^2^ to evaluate the performance in batch effect correction [22, 48]. We evaluate and visualize all the above metrics based on the package scib-metrics [22]. The overall score is calculated as a weighted average of all metrics on bio-conservation (40% weights), modality integration (30% weights), and batch correction (30% weights).

In our experiments, we set the batch size to 128, the maximum epoch number to 200, and the dimensions of both the shared latent component (*d*_*u*_) and modality-specific component (*d*_*s*_) to 20. Parameters are optimized via Adam optimizer and the learning rate is started from 0.001 with a cosine annealing scheduler. 10% of the data was used as a validation set for early stopping during training based on the total loss on validation set. For scATAC-seq data, peak count signals are converted to gene-level gene activity scores using EpiScanpy. For datasets integrating two modalities, we select highly variable genes that are measured in both scRNA-seq and scATAC-seq (based on gene activity scores) as input features, and do not apply masked loss. For tri-modal integration, we use highly variable genes from scRNA-seq and scATAC-seq, as well as all genes with nonzero measurements in scProtein as input features, and apply masked loss according to the available gene list associated with each modality. Detailed hyperparamters setting is shown in Supplementary Table S1.

## Supplementary information

Supplementary materials are available online.

## Acknowledgements

This work is supported by Shanghai Artificial Intelligence Laboratory.

## Declarations

- Conflict of interest: No conflict of interest.
- Ethics approval and consent to participate: Not applicable.
- Consent for publication: Not applicable.
- Data availability: Benchmark single-cell data used are available at https://scglue.readthedocs.io/en/latest/data.html and https://cellxgene.cziscience.com/collections/ae1420fe-6630-46ed-8b3d-cc6056a66467, paired 10x multiome data are available at https://www.10xgenomics.com/datasets, mouse primary motor cortex merFISH omics data are available at https://cellxgene.cziscience.com/collections/9132fae8-bdfe-480f-9e45-45bc77f320b3, mouse frontal cortex data are available at https://drive.google.com/drive/folders/1smPbTfkkdPvk0hXzIZrXk55aEbnXPVl?usp=sharing, 10x visum data are available at https://satijalab.org/seurat/articles/spatial_vignette, mouse methylation data are available at https://brainome.ucsd.edu/annoj/brain_single_nuclei/, the multi-species scRNA data are available at https://assets.nemoarchive.org/dat-zkgd0it.
- Code availability: The codes are available at https://github.com/sjl-sjtu/scMRDR.

## Supplementary Materials

### Supplementary Notes

#### SN1 Benchmark experiments settings and evaluation metrics Experiments settings

The species, tissues, sample sizes, modalities, and references (the original sources) of the single-cell datasets used in the experiments are shown in Supplementary Table S1. In our experiments, we set the batch size to 128, the epoch number to 200, and the dimensions of both the shared latent component (*d*_*u*_) and modality-specific component (*d*_*s*_) to 20. Parameters are optimized via Adam optimizer and the learning rate is started from 0.001 with a cosine annealing scheduler. Other hyperparameter settings are summarized in Table Supplementary Table S1. 10% of the data was used as a validation set for early stopping during training based on the total loss on validation set. For scATAC-seq data, peak count signals are converted to gene-level gene activity scores using EpiScanpy. For datasets integrating two modalities, we select highly variable genes that are measured in both scRNA-seq and scATAC-seq (based on gene activity scores) as input features, and do not apply masked loss. For tri-modal integration, we use highly variable genes from scRNA-seq and scATAC-seq, as well as all genes with nonzero measurements in scProtein as input features, and apply masked loss according to the available gene list associated with each modality.

Competing methods are used with their respective default settings. Specifically, Seurat, Harmony, and scVI use gene activity scores for scATAC-seq, while all other methods operate on the original peak counts.

We run all empirical experiments on a single NVIDIA RTX 4080 GPU. The runtime of our method for 200 epochs is also reported in Supplementary Table S1.

#### Evaluation metrics

- F1 isolated label scores. The optimal F1 score by optimizing the cluster assignment of the isolated label using the F1 score across Louvain clustering (resolutions 0.1–2, step 0.1). The metric is averaged across all isolated cell-type labels.
- Silhouette scores. The global silhouette width measures (ASW) between isolated and non-isolated labels on the PCA embedding, scaled to [0, 1]. ASW measures how similar a data point is to its own cluster compared to other clusters, with higher ASW indicating more compact and well-separated clusters. For the bio-conservation, ASW was computed and averaged across cell identity labels; while for modality or batch integration, ASW was computed and averaged across batch or modality labels, and then subtracted from 1 to ensure higher scores indicate better integration.
- Kmeans NMI. Normalized Mutual Information (NMI) measures the similarity between clustering results and known labels, accounting for label permutations. We compute NMI between KMeans clusters and ground truth labels, with scores ranging from 0 (no overlap) to 1 (perfect match).
- Kmeans ARI. Adjusted Rand Index (ARI) evaluates clustering accuracy by considering both agreements and disagreements between predicted and true labels, adjusted for chance. We compare KMeans clusters with ground truth labels, where 0 indicates random labeling and 1 indicates perfect agreement.
- Graph LISI. Graph LISI is an extension of the original LISI metric that is computed from neighborhood lists per node from integrated kNN graphs. Instead of relying on a fixed number of nearest neighbors, Graph LISI computes shortest-path distances on the integrated graph to consistently define neighborhoods for each cell, providing a stable diversity score even when the underlying graph has variable connectivity. The resulting scores are rescaled to a 0–1 range, where higher values indicate better batch or modality integration (iLISI) or better cell-type separation (cLISI).
- PCR. Principal Component Regression (PCR) is used to quantify batch effects by assessing how much variance in the data can be attributed to batch variables or modality differences, which is computed by multiplying the variance explained by each principal component (PC) with the *R*^2^ value from regressing the batch on that PC and then summing over all PCs.
- kBET. k-nearest neighbor Batch Effect Test (kBET) assesses data mixing by checking whether the local batch label distribution in each cell’s neighborhood matches the global distribution. It reports a rejection rate across tested neighborhoods, where a lower rate indicates better batch mixing.
- Graph connectivity. Graph connectivity, ranging from 0 to 1, measures how well cells of the same identity are connected within the integrated kNN graph. For each cell type, it computes the fraction of cells in the largest connected component of that type’s subgraph.

### Supplementary Tables

**Supplementary Table S1.**
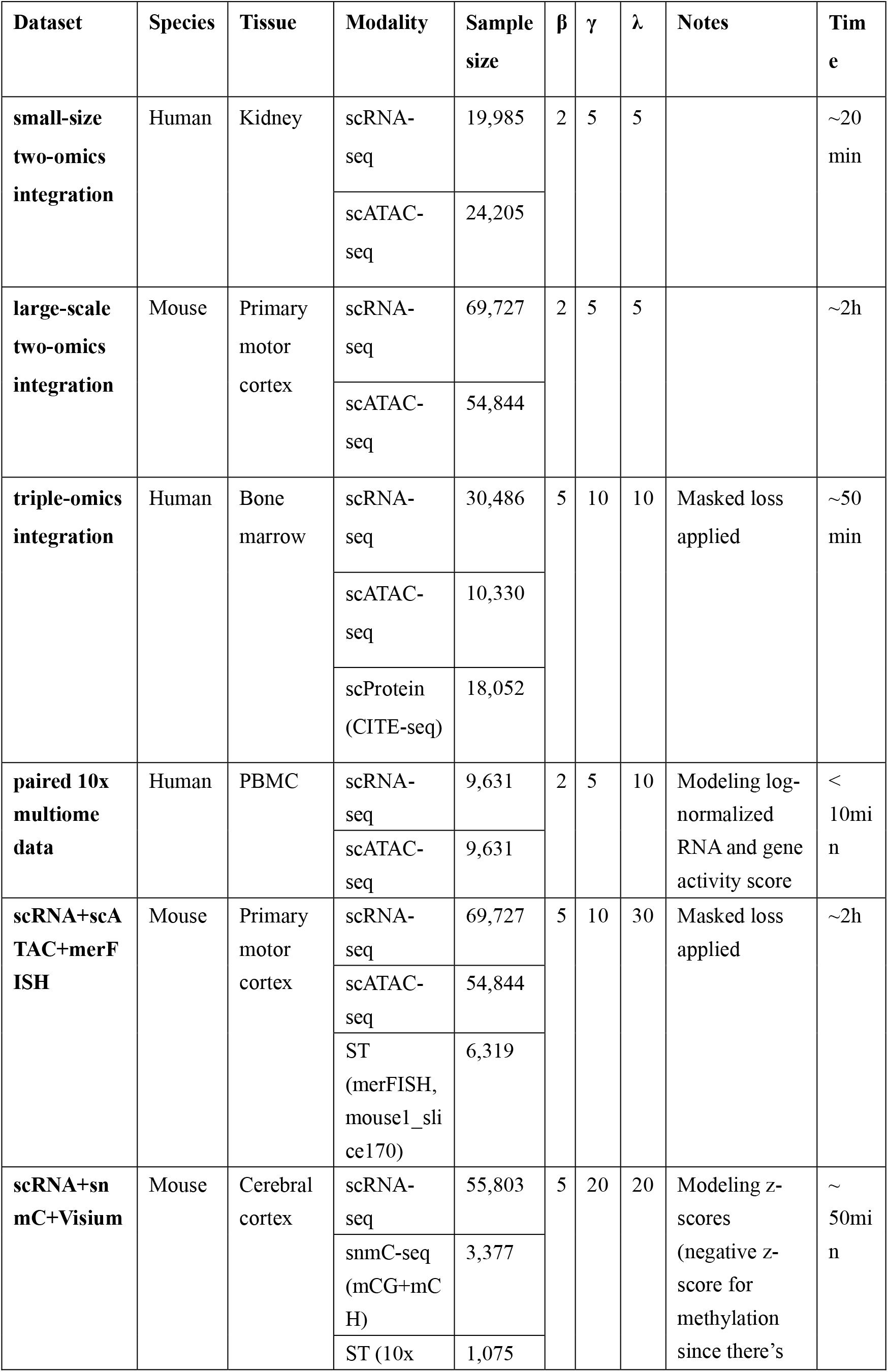
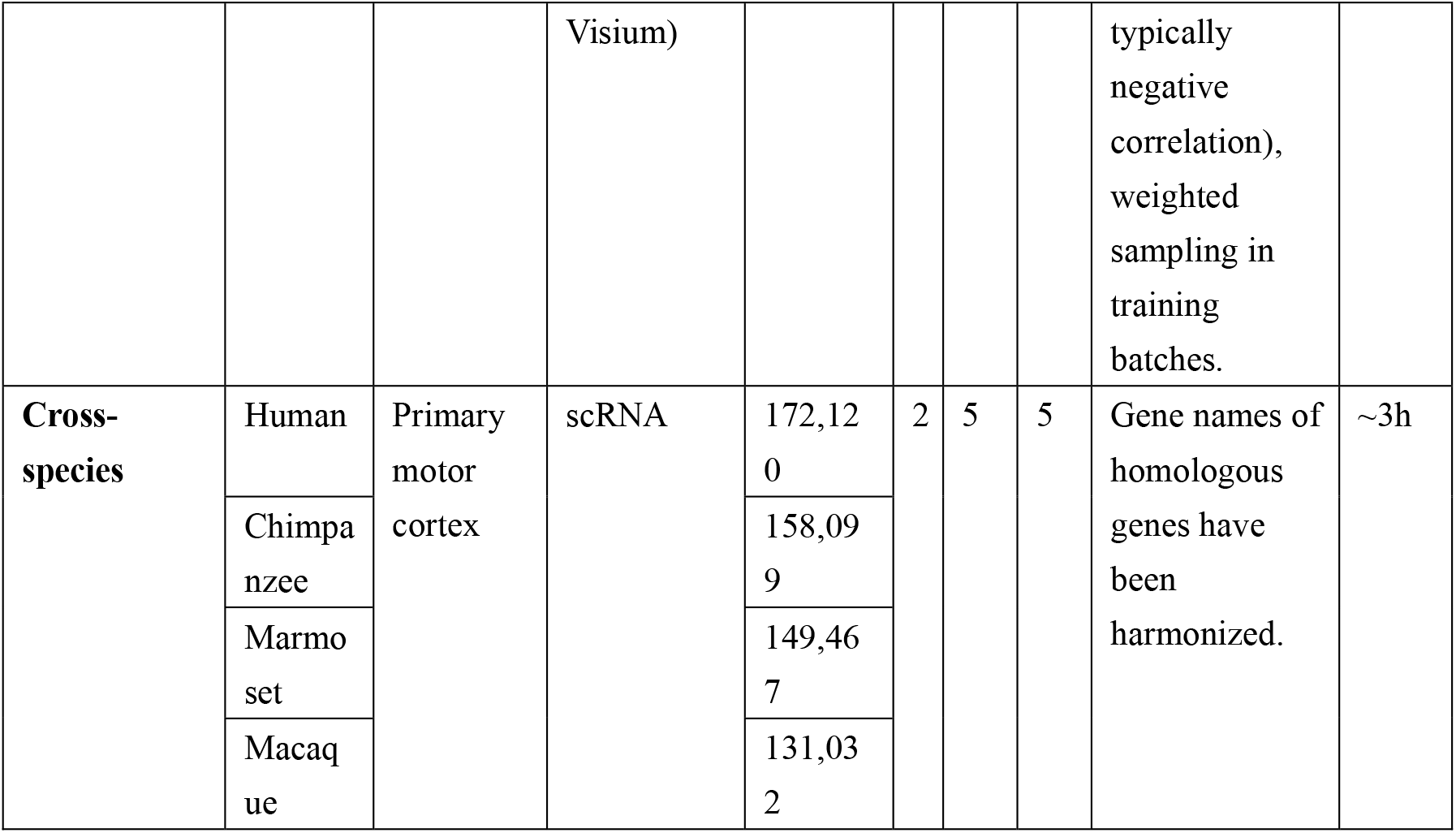
Experiments details.

**Supplementary Table S2.** Performance in ablation studies and sensitivity analysis, where unscaled scIB aggregate scores are reported.

| $\beta$ | $\gamma$ | $\lambda$ | Total Score | Batch-correct | Bio-conserve | Modal-integrate | Notes |
| --- | --- | --- | --- | --- | --- | --- | --- |
| 2 | 5 | 5 | 0.66 | 0.52 | 0.74 | 0.70 |  |
| 2 | 7 | 5 | 0.65 | 0.54 | 0.69 | 0.72 |  |
| 2 | 5 | 3 | 0.65 | 0.55 | 0.69 | 0.71 |  |
| 2 | 2 | 2 | 0.65 | 0.50 | 0.72 | 0.70 |  |
| 2 | 5 | 7 | 0.65 | 0.49 | 0.70 | 0.73 |  |
| 2 | 3 | 5 | 0.65 | 0.47 | 0.71 | 0.74 |  |
| 2 | 1 | 5 | 0.64 | 0.49 | 0.68 | 0.75 |  |
| 3 | 5 | 5 | 0.64 | 0.48 | 0.71 | 0.71 |  |
| 4 | 5 | 5 | 0.64 | 0.47 | 0.71 | 0.72 |  |
| 2 | 10 | 5 | 0.64 | 0.52 | 0.68 | 0.71 |  |
| 1 | 5 | 5 | 0.62 | 0.50 | 0.67 | 0.68 | $\beta = 1$ |
| 2 | 5 | 0 | 0.61 | 0.53 | 0.69 | 0.60 | $\lambda = 0$ |
| 2 | 0 | 5 | 0.61 | 0.51 | 0.60 | 0.72 | $\gamma = 0$ |
| 1 | 0 | 0 | 0.59 | 0.48 | 0.63 | 0.66 | **Baseline** |

**Supplementary Table S3.** Ablation study in triple-omics data, where unscaled scIB aggregate scores are reported.

|  | Overall Score | Batch Correct | Bio Conserve | Modal Integrate |
| --- | --- | --- | --- | --- |
| **ours** | 0.58 | 0.39 | 0.69 | 0.61 |
| $\beta = 1$ | 0.51 | 0.39 | 0.58 | 0.52 |
| $\lambda = 0$ | 0.48 | 0.32 | 0.60 | 0.49 |
| $\gamma = 0$ | 0.53 | 0.40 | 0.58 | 0.58 |
| **w/o truncated loss** | 0.44 | 0.33 | 0.58 | 0.37 |

### Supplementary Figures

**Supplementary Fig. S1.**
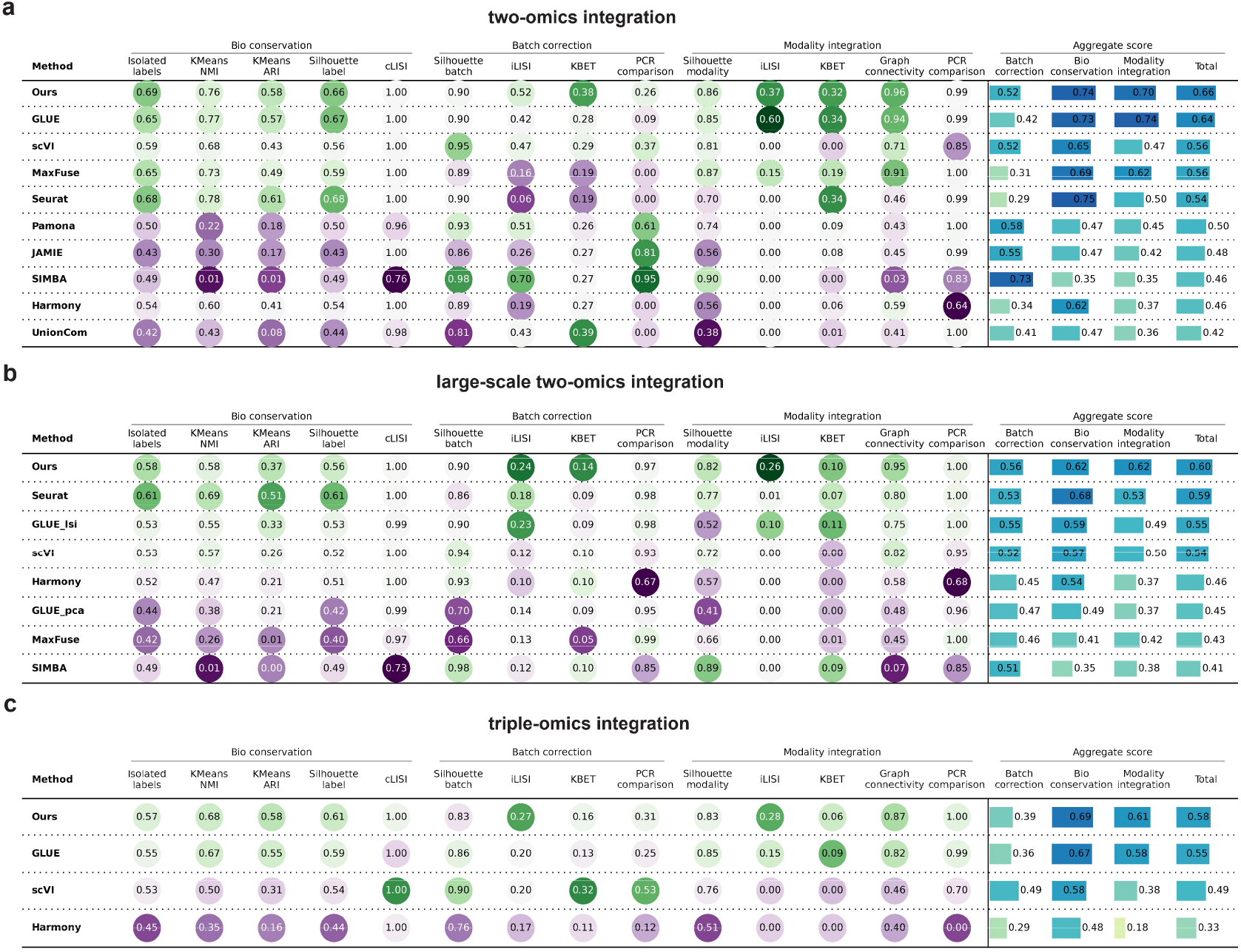
Details on the unscaled scIB evaluation metrics in two-omics integration task (a), large-scale two-omics integration task (b), and triple-omics integration task (c).

**Supplementary Fig. S2.**
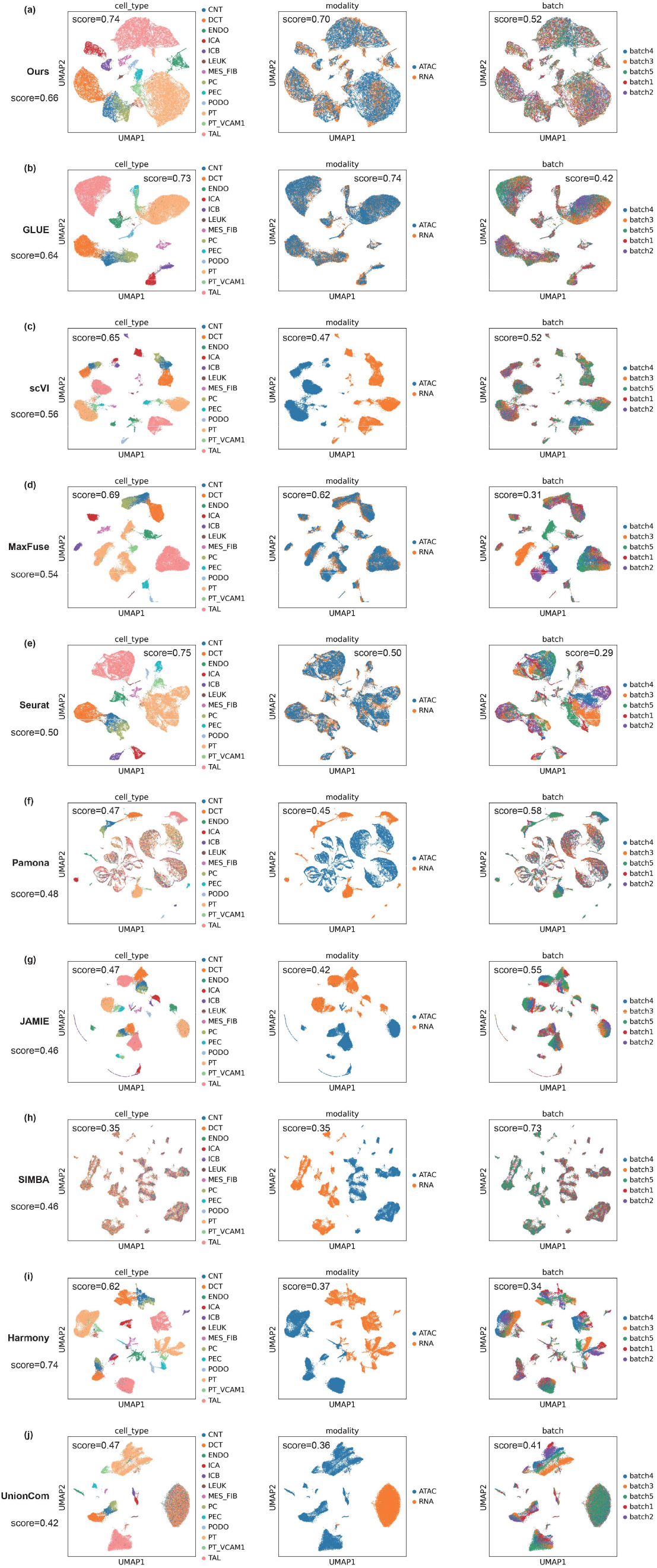
UMAP visualization of the unified embeddings obtained by different methods in small-scale two-omics data integration task.

**Supplementary Fig. S3.**
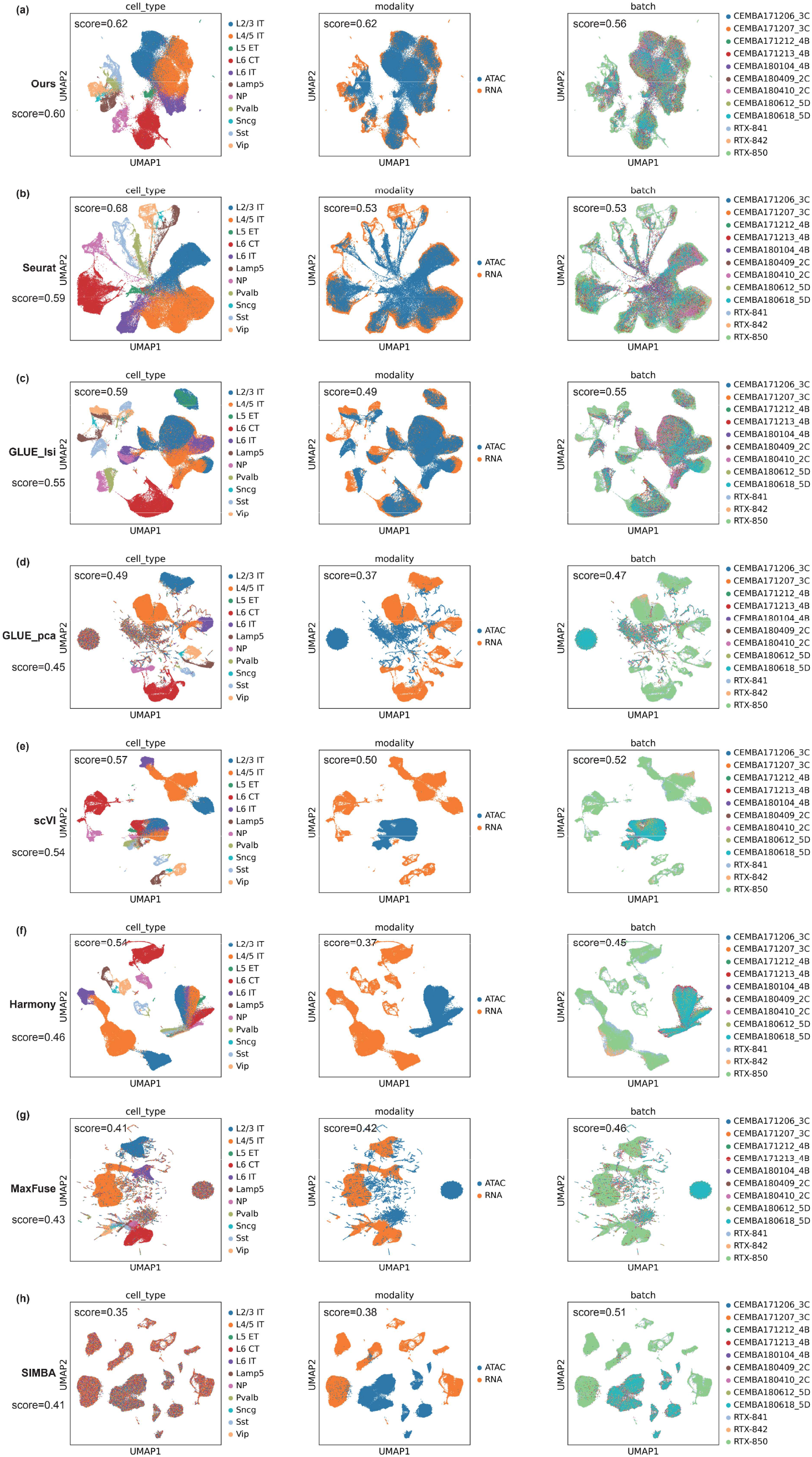
UMAP visualization of the unified embeddings obtained by different methods in large-scale two-omics data integration task.

**Supplementary Fig. S4.**
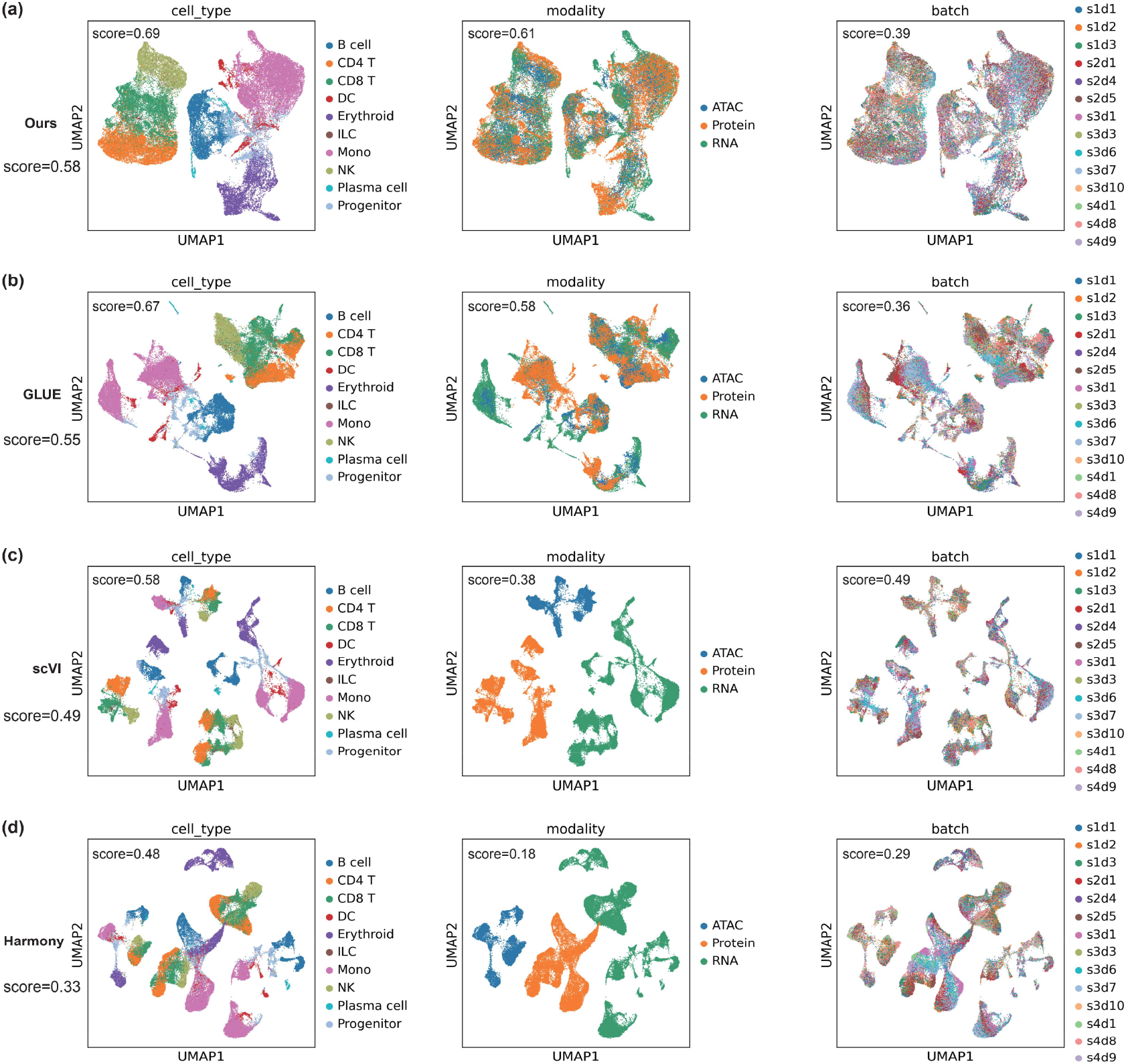
UMAP visualization of the unified embeddings obtained by different methods in triple-omics data integration task.

**Supplementary Fig. S5.**
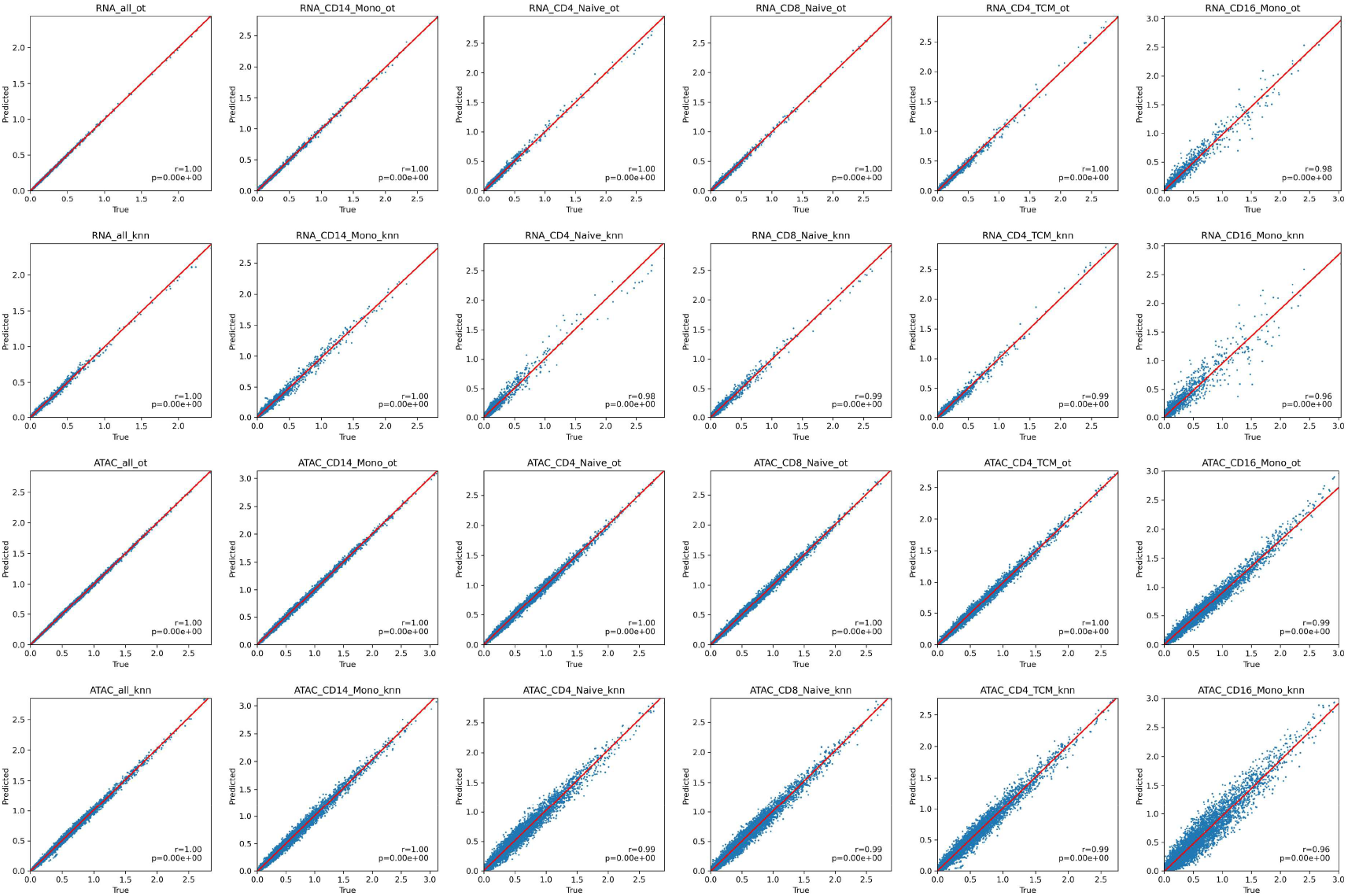
Correlation between the true and predicted mean RNA expressions / ATAC activity scores of different genes within all cells and cells from the top 5 cell types with the highest quant.

**Supplementary Fig. S6.**
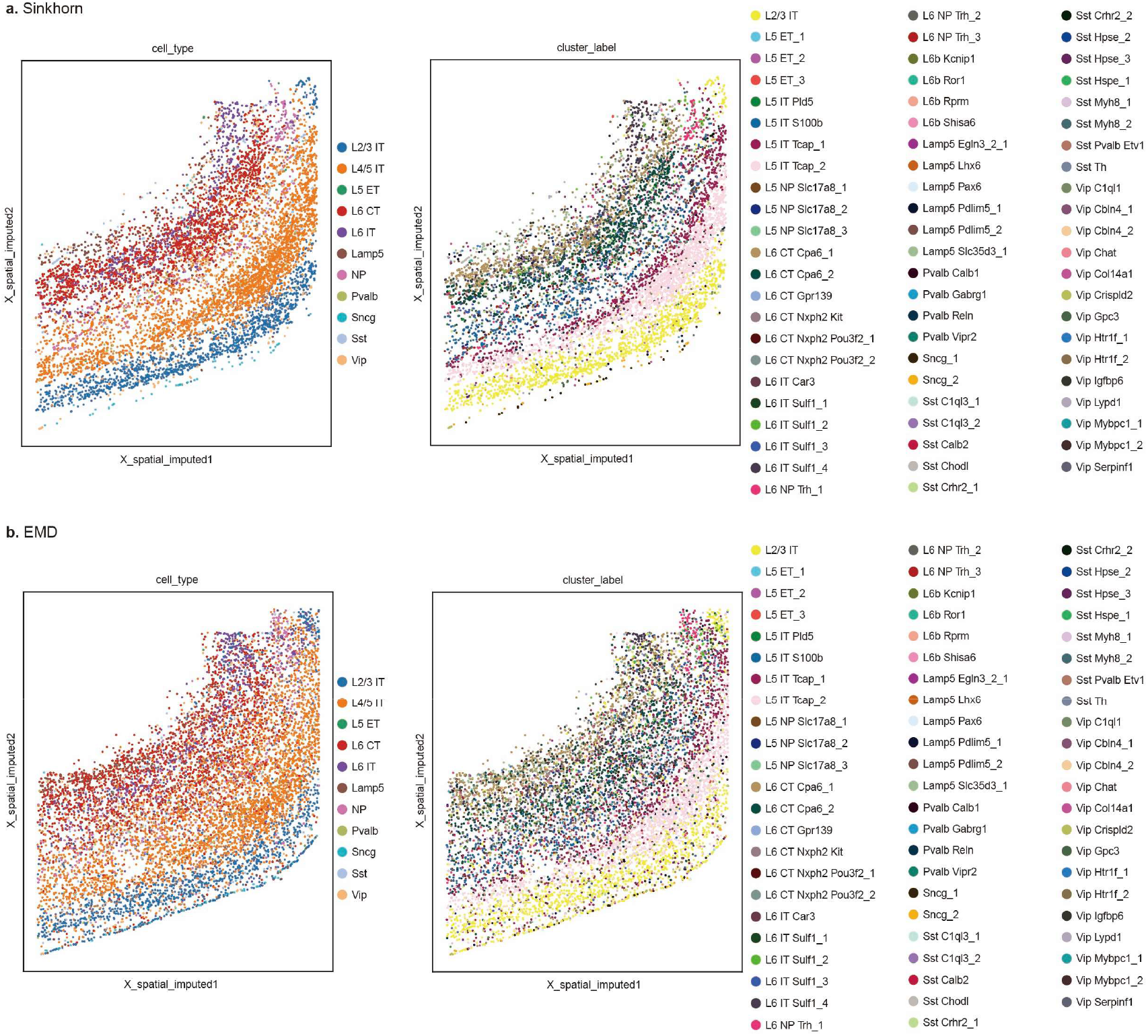
Spatial visualization of mouse primary motor cortex scRNA data, where the cell type and subtype labels are provided by the original study, and the spatial locations are imputed by scMRDR. Two OT variants - Sinkhorn (a) and EMD (b) algorithms - are used in imputing spatial locations.

**Supplementary Fig. S7.**
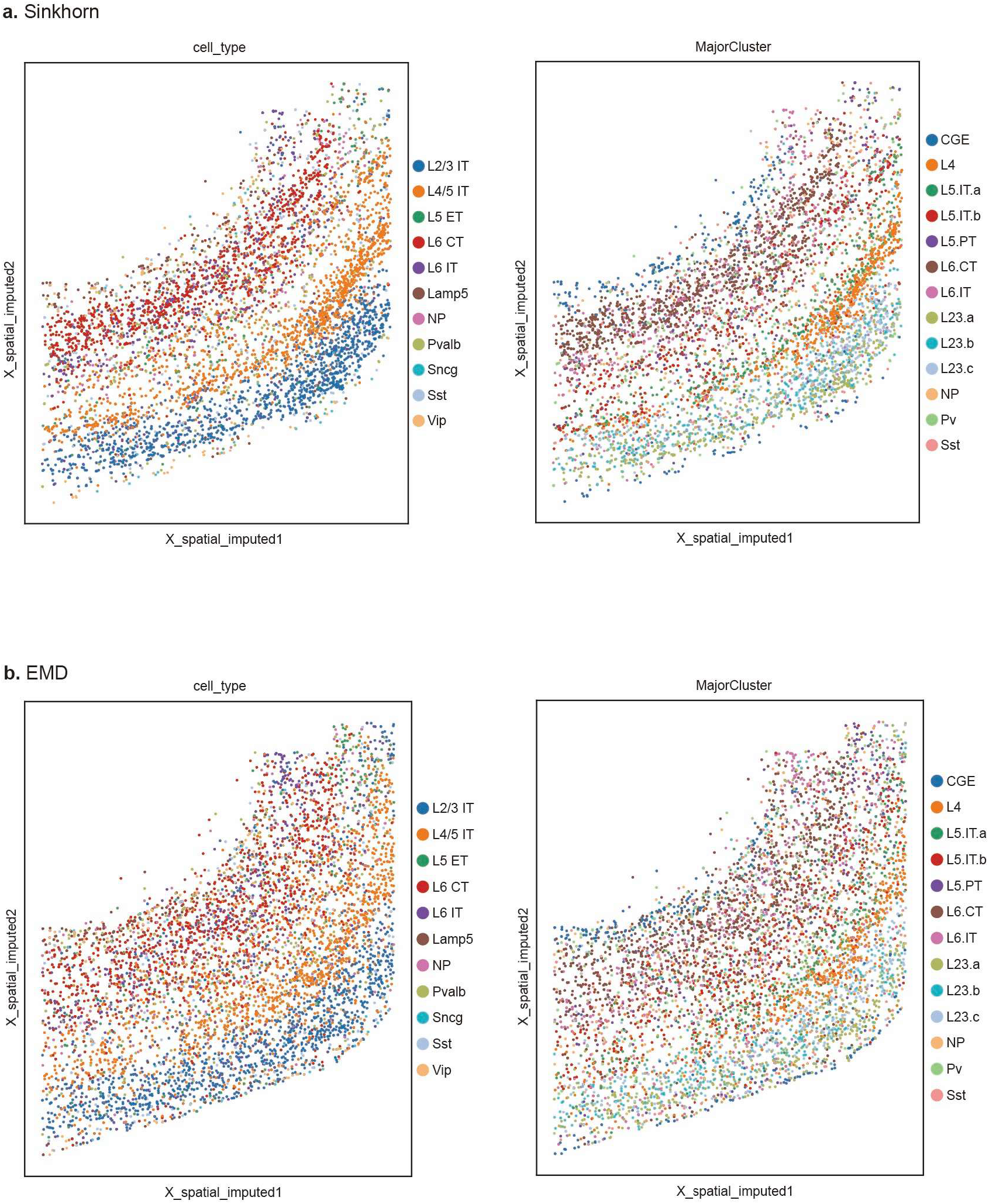
Spatial visualization of mouse primary motor cortex scATAC data, where the cell type and subtype labels are provided by the original study, and the spatial locations are imputed by scMRDR. Two OT variants - Sinkhorn (a) and EMD (b) algorithms - are used in imputing spatial locations.

**Supplementary Fig. S8.**
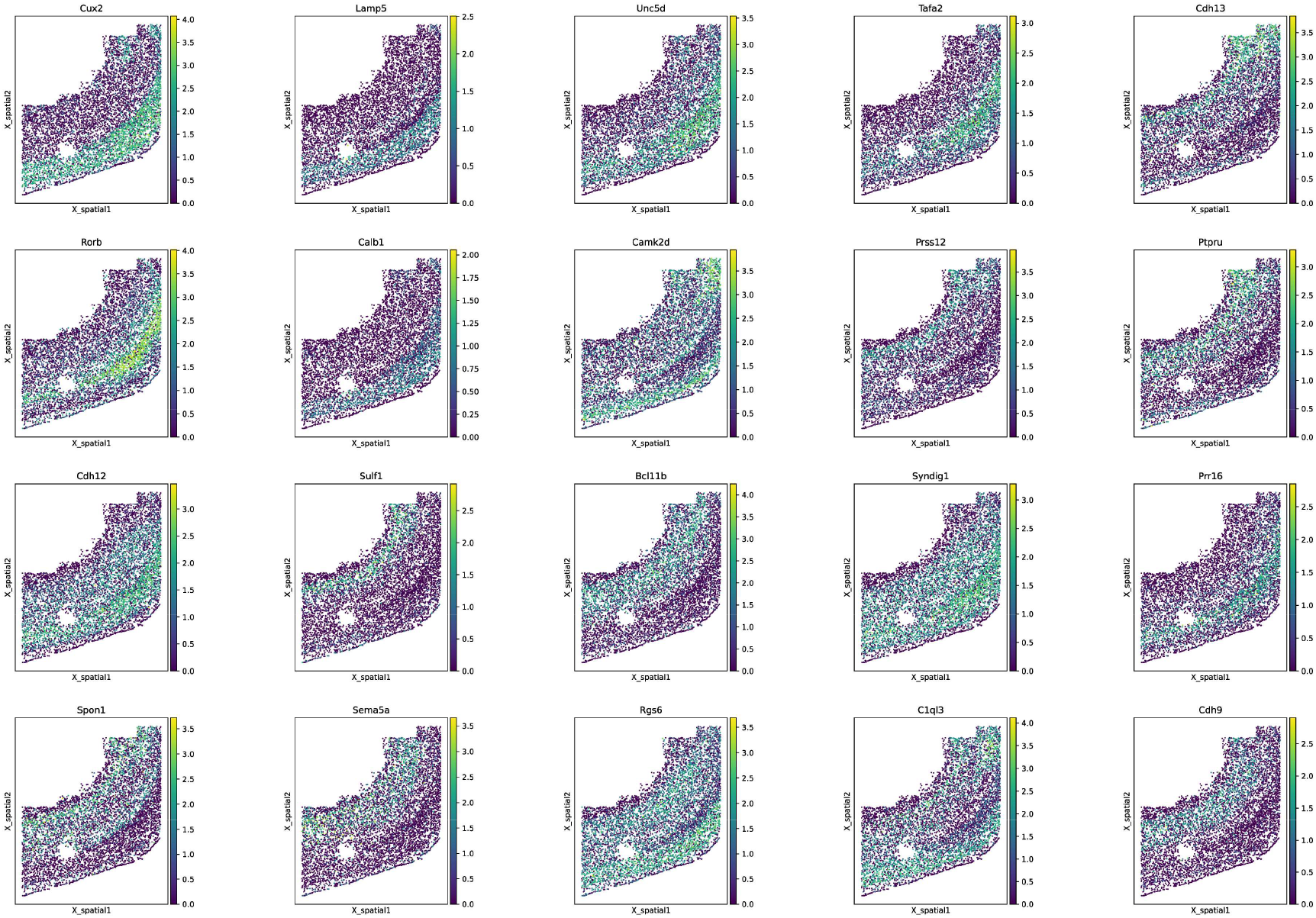
Top 20 SVGs detected via SPARKX on the original merFISH ST data on mouse primary motor cortex.

**Supplementary Fig. S9.**
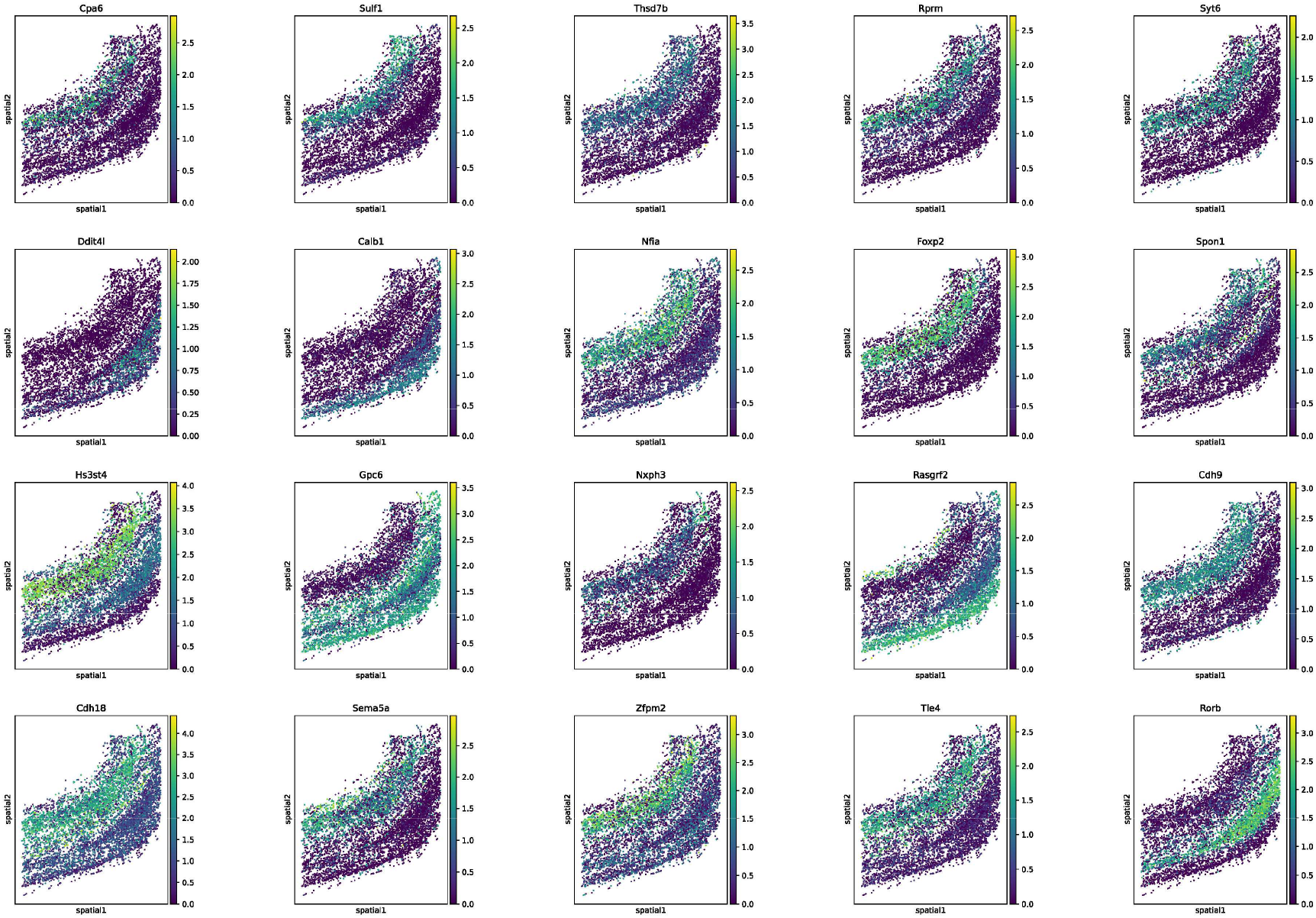
Top 20 SVGs detected via SPARKX on the scRNA data with imputed spatial locations on mouse primary motor cortex.

**Supplementary Fig. S10.**
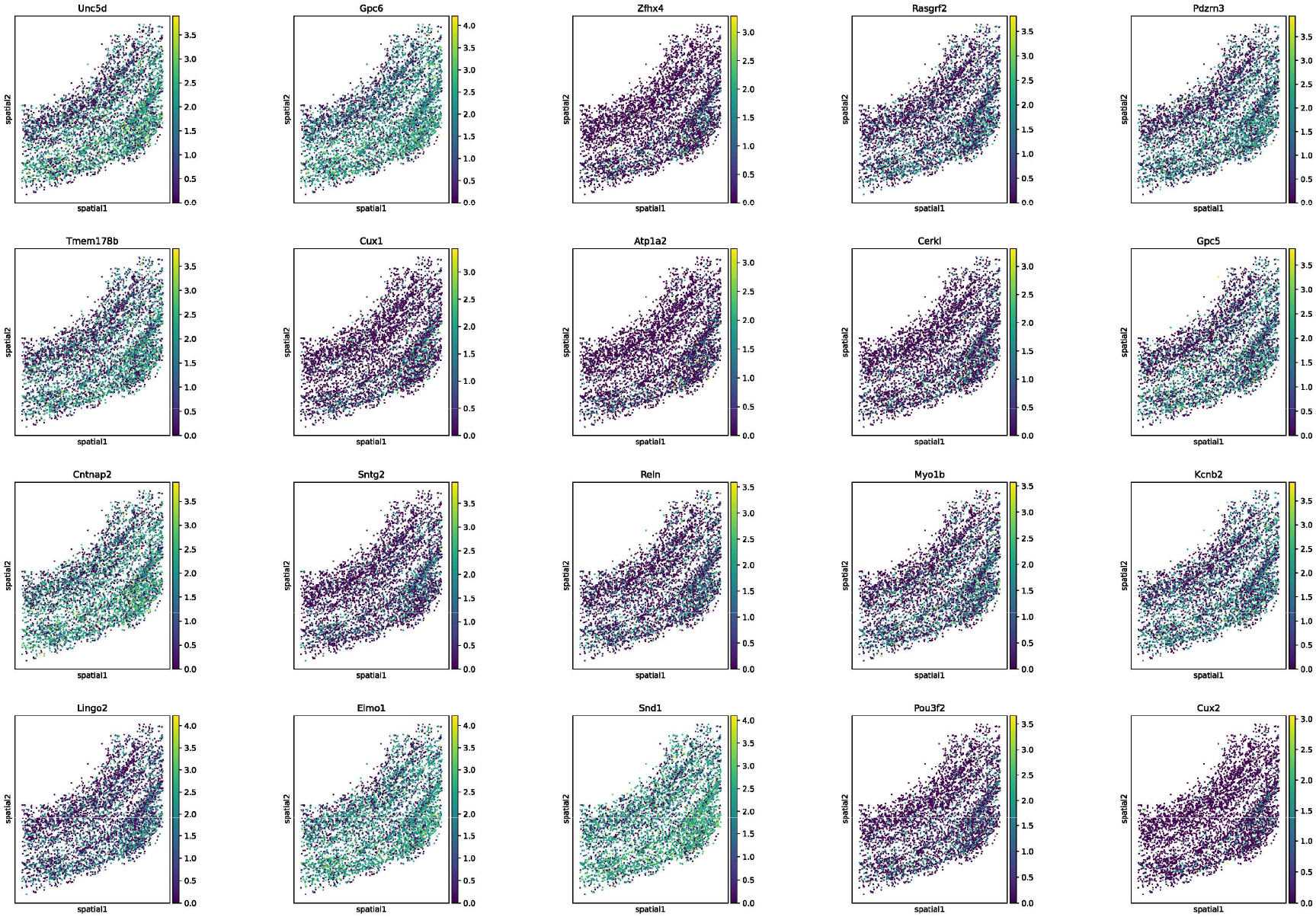
Top 20 SVGs detected via SPARKX on the gene activity score calculated from scATAC data with imputed spatial locations on mouse primary motor cortex.

**Supplementary Fig. S11.**
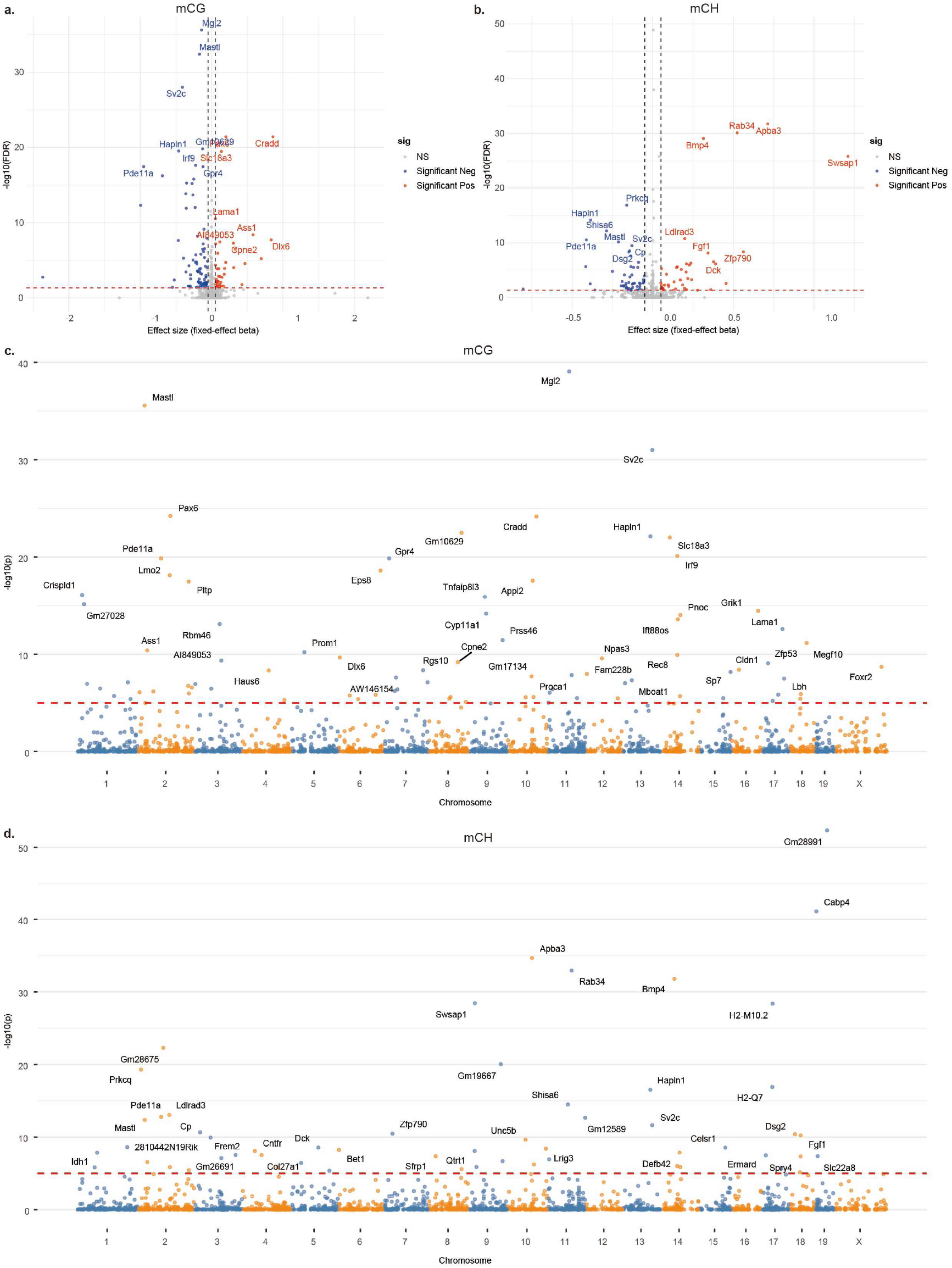
Gene methylation’s effects on expression tested via Spatial Generalized Additive Model (SGAM) accounting for spatial autocorrelation and cell-type heterogeneity, where cell types are incorporated as a fixed effect. Volcano plots show effect size (beta) and BH-adjusted FDR for mCG (a) and mCH (b) sites. Manhattan plots show the effect significance for mCG (c) and mCH (d), where the red dashed line indicates the significance threshold 1e-5 (since thousands of genes are included).

**Supplementary Fig. S12.**
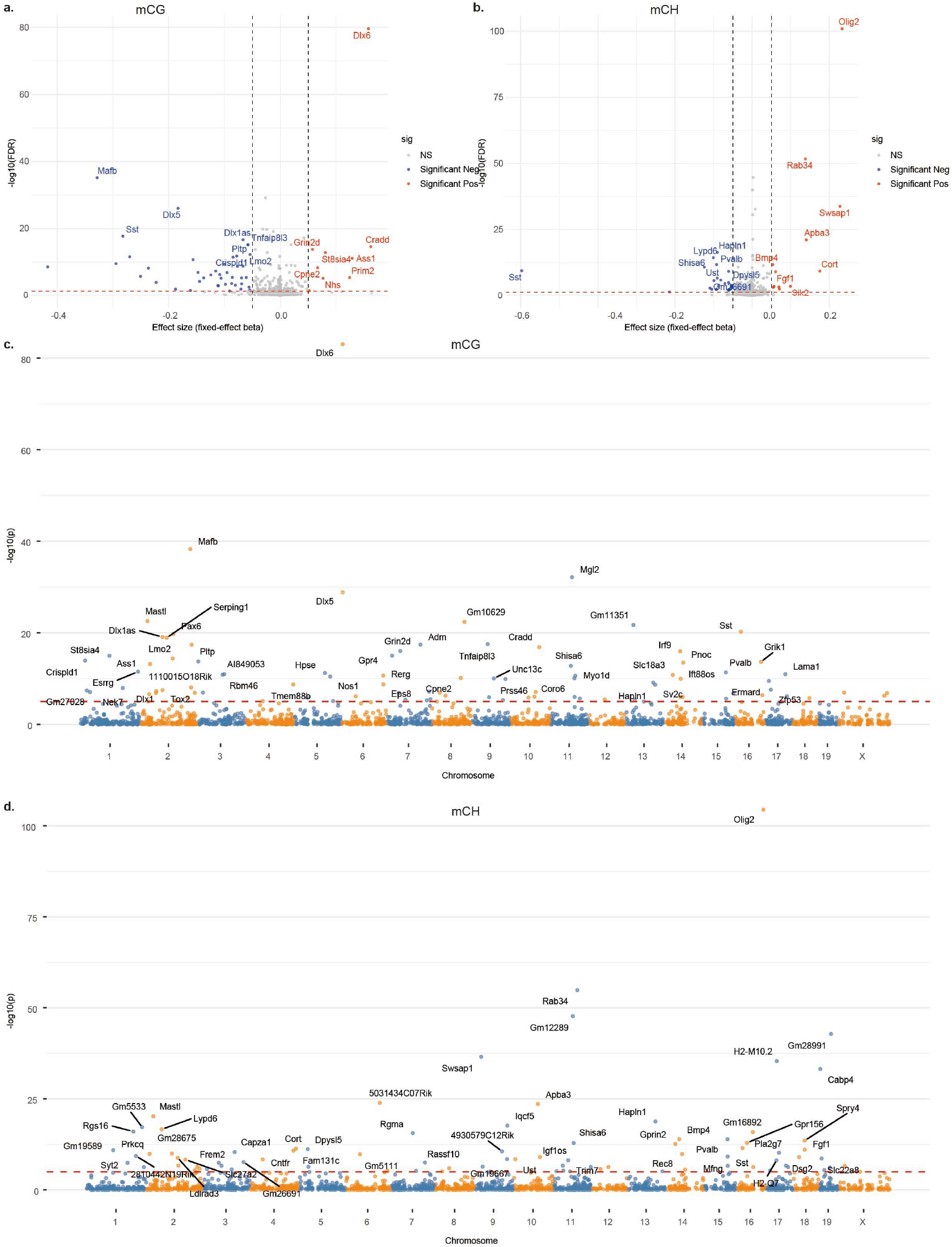
Gene methylation’s effects on expression tested via Spatial Generalized Additive Mixed Model (SGAMM) accounting for spatial autocorrelation and cell-type heterogeneity, where cell types are incorporated as a random effect. Volcano plots show effect size (beta) and BH-adjusted FDR for mCG (a) and mCH (b) sites. Manhattan plots show the effect significance for mCG (c) and mCH (d), where the red dashed line indicates the significance threshold 1e-5 (since thousands of genes are included).

**Supplementary Fig. S13.**
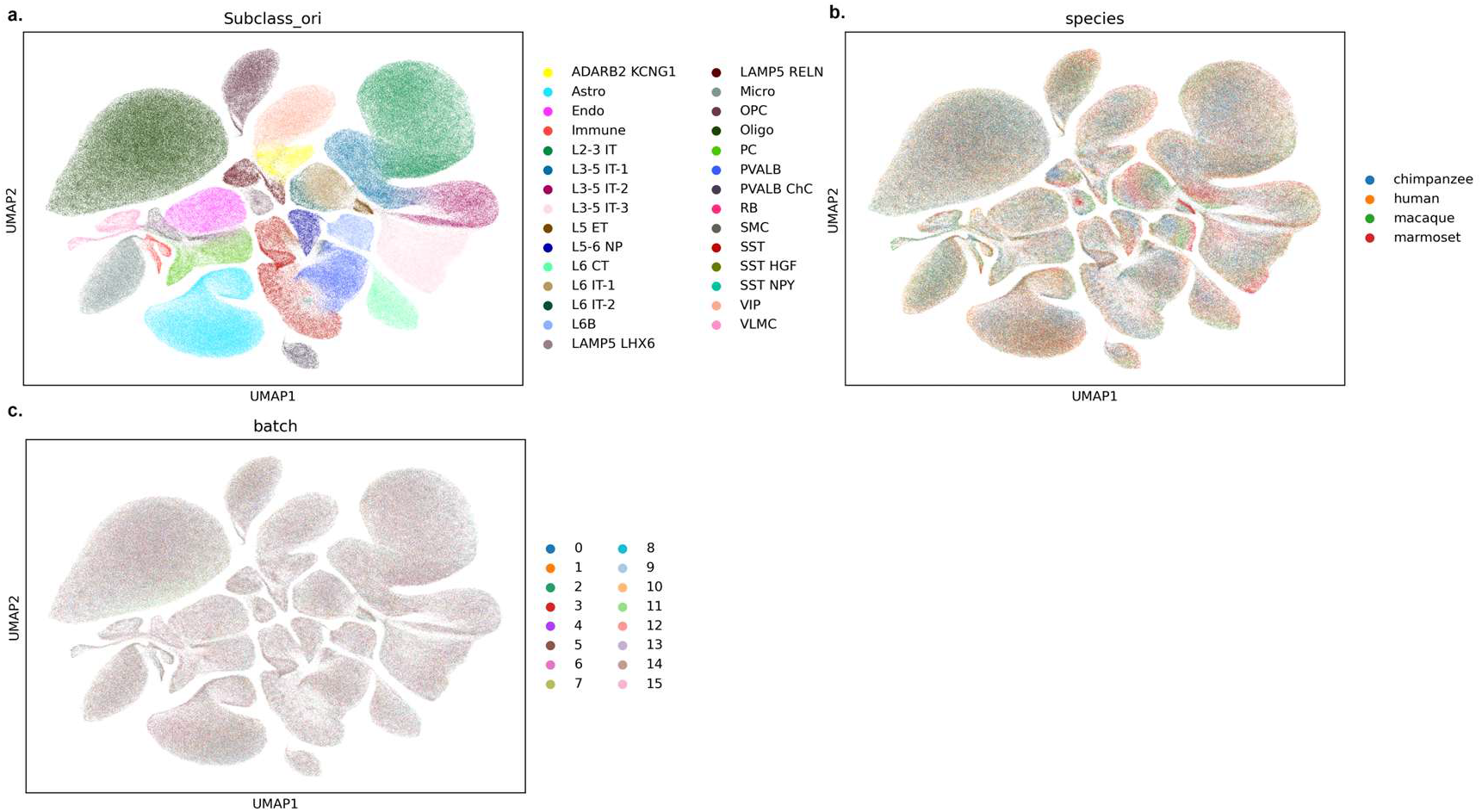
UMAP visualization of cross-species scRNA integration of primary motor cortex cells, where cells are colored by their subclass labels (a), species (b), and batch information (c).

